# Human Hepatocyte Nuclear Factor 4-α encodes isoforms with distinct transcriptional functions

**DOI:** 10.1101/585604

**Authors:** Élie Lambert, Jean-Philippe Babeu, Joël Simoneau, Dominique Lévesque, Émilie Jolibois, Michelle Scott, François Boudreau, François-Michel Boisvert

## Abstract

HNF4α is a nuclear receptor produced as 12 isoforms from two promoters by alternative splicing. In order to characterize the transcriptional capacities of all 12 HNF4α isoforms, stable lines expressing each isoform were generated. The entire transcriptome associated with each isoform was analyzed as well as their respective interacting proteome. Major differences were noted in the transcriptional function of these isoforms. The α1 and α2 isoforms were the most potent regulators of gene expression while the α3 isoform exhibited significantly reduced activity. The α4, α5 and α6 isoforms, which use an alternative first exon, were characterized for the first time, and showed a greatly reduced transcriptional potential with an inability to recognize the consensus response element of HNF4α. Several transcription factors and coregulators were identified as potential specific partners for certain HNF4α isoforms. An analysis integrating the vast amount of omics data enabled the identification of transcriptional regulatory mechanisms specific to certain HNF4α isoforms, hence demonstrating the importance of considering all isoforms given their seemingly diverse functions.

## INTRODUCTION

Nuclear receptors (NR) represent a class of transcription factors that encompasses 48 proteins in humans (Zhang et al., 2004). The nomenclature surrounding the superfamily of NR, based on their phylogeny, consists of six subfamilies comprised of several groups (Nuclear Receptors Nomenclature Committee, 1999). NR share a structural organization of five to six distinct regions designated from A to F (Robinson-Rechavi et al., 2003). The A/B region, located at the N-terminal end, is highly variable between the various NR. It typically contains an AF-1 transactivation region (Activation Function) which is active independently of ligand binding and allows the interaction of the receptor with various coregulators and other transcription factors (Lavery and McEwan, 2005). The C domain, or DNA-binding domain (DBD), is the most conserved among NR. It allows the recognition of specific DNA response elements via two cysteine-rich zinc finger motifs (C-X2-C-X13-C-X2-C and C-X5-C-X9-C-X2-C) (Robinson-Rechavi et al., 2003). These response elements are composed of repeated or inverted hexameric DNA sequences and separated by linkers varying from one to five nucleotides in length (Khorasanizadeh and Rastinejad, 2001). The D region, also called hinge region, is less conserved and its main function is to facilitate free rotation between the DBD and the ligand binding domain (LBD). A nuclear localization signal (NLS) contained in this region participates in the regulation of the subcellular distribution of NR (Germain et al., 2006). The E domain, or ligand binding domain (LBD), consists of a hydrophobic pouch for binding a multitude of small lipophilic molecules such as steroid hormones, phospholipids, fatty acids and xenobiotics (Pawlak et al., 2012). Some NR for which no ligand has yet been identified are considered orphan receptors (Giguere, 1999). A second AF-2 activator region is located in the LBD. In contrast to the AF-1 region, the activity of the AF-2 region is dependent on ligand binding to LBD. This induces a conformational change in the LBD, generating a pouch that can interact with the LXXLL motif present on a host of transcriptional coactivators (Heery et al., 1997). The LBD, like the DBD, contains an important interface for receptor dimerization. Finally, at the C-terminus of the NR is the F domain. Because of its highly variable sequence, the exact function of the F domain remains to be established and several NR have no F domain (Patel and Skafar, 2015). Nevertheless, the deletion of this domain in those receptors bearing the domain have revealed its importance in certain instances in connection with various functions such as dimerization, activation and interaction with different coregulators (Patel and Skafar, 2015).

HNF4α (Hepatocyte Nuclear Factor 4 alpha) (also referred to as NR2A1) is a transcription factor of the nuclear receptor family that was initially identified as a regulator of liver-specific gene expression (Costa et al., 1989; Sladek et al., 1990). Since its initial discovery in the liver, HNF4α has also been detected in the kidneys, pancreas, stomach, small intestine and colon (Tanaka et al., 2006). HNF4α is crucial for the development and maintenance of hepatocyte function, including lipid homeostasis, transport and metabolism, as well as the detoxification of xenobiotics (Hayhurst et al., 2001; Wortham et al., 2007; Yin et al., 2011). Additional functions for HNF4α in the gut and pancreas have also emerged (Drewes et al., 1996; Eeckhoute et al., 2003; Wang et al., 2000). In contrast to other types of NR, HNF4α is constitutively localized in the nucleus and does not require binding of a ligand in order to homodimerize and interact with the response elements of its target genes (Yuan et al., 2009). HNF4α has long been considered an orphan nuclear receptor, although crystallography of the LBD initially revealed the presence of fatty acids of various compositions bound at the level of the ligand binding pocket of HNF4α (Dhe-Paganon et al., 2002). Subsequent studies have identified linoleic acid, a long chain polyunsaturated fatty acid (C18: 2ω6), as the molecule preferentially binding to its LBD (Yuan et al., 2009); however, this binding is reversible and does not modulate the transcriptional activity of HNF4α. The nature of the HNF4α ligand therefore remains controversial, since linoleic acid is endogenously present and does not appear necessary for receptor activity, unlike the typical mode of action of NR requiring binding to their ligand. HNF4α recognizes DR1 sites (Direct Repeat 1), consisting of two repeated hexameric half-sites separated by a nucleotide, typically an adenosine (Fang et al., 2012). HNF4α also recognizes direct repeats separated by two nucleotides (DR2), but with lower specificity (Jiang and Sladek, 1997). The consensus sequence of its half-sites, AGGTCA, is shared by most nonsteroidal NR (Fang et al., 2012). HNF4α is considered to be an exclusive homodimer, this form being stably found in solution and is necessary to bind DNA (Jiang et al., 1995). However, both RXRα/β/γ and RARα nuclear receptors, known for their ability to form heterodimers with several NR, do not assemble into heterodimers with HNF4α (Jiang et al., 1995; Lee and Privalsky, 2005).

Alternative splicing is a major source of cellular protein diversity. The estimated percentage of human gene products undergoing alternative spicing has been proposed to be as high as 95% of multiexon genes, although it is still unclear how many of these splicing variants are actually expressed or functional (Pan et al., 2008). However, very few of these alternative protein isoforms have well-characterized cellular functions, given that studies on these proteins have either mostly concentrated on a single isoform, or do not specify which isoform was under study, hence leading to discrepancies or contradictions in protein function (Kelemen et al., 2013). In addition, a large number of alternatively spliced transcripts studied for protein-protein interaction by yeast two-hybrid assay were shown to display significant differences between reference and alternative isoforms, with many alternative isoforms interacting with different protein partners (Yang et al., 2016). These differences in protein complexes underline the importance of considering each protein isoform in order to understand its unique role(s).

In the present study, the transcriptional functions of the 12 annotated isoforms of HNF4α were specifically characterized (Babeu and Boudreau, 2014; Sladek and Seidel, 2001) by generating stable lines expressing each HNF4α isoform in HCT 116 cells. The entire transcriptome associated with each isoform was analyzed by RNA sequencing, as well as their respective proteome by a BioID approach coupled to quantitative mass spectrometry. This analysis integrating the vast amount of transcriptomic and proteomic data enabled the identification of transcriptional regulatory mechanisms specific to each isoform, demonstrating the importance of considering all isoforms which can exhibit distinct functions.

## RESULTS

### New classification of HNF4α isoforms

An important issue arising from the existence of the HNF4α isoforms is the lack of detail and uniformity in their descriptions in the literature and databases. Indeed, the main protein databases such as RefSeq (O’Leary et al., 2016), Uniprot (UniProt, 2019) and Ensembl (Zerbino et al., 2018) use different numbering and identification strategies for protein isoforms, and studies on proteins in the literature often do not specify which isoform is under consideration (Supplementary Table 1). Since the initial identification of HNF4α in the liver, a total of 12 HNF4α isoforms have been reported or predicted in the literature encompassing protein isoforms with distinct N- and C-termini regions (Babeu and Boudreau, 2014; Huang et al., 2009; Sladek and Seidel, 2001). A first classification of the different isoforms separates the different proteins depending on promoter usage (termed P1 and P2, (Boj et al., 2001; Thomas et al., 2001)) generating differences in the N-terminus which will include either exon 3 (isoforms α1-3) or exon 4 (isoforms α4-6) for isoforms transcribed from the P1 promoter, or exon 1 (isoforms α7-9) or exon 2 (isoforms α10-12) for isoforms transcribed from the P2 promoter (Figure 1). Interestingly, only the N-termini from isoforms α1-3 include the activation function 1 domain (AF-1 domain), whereas it has not been identified in the the other isoforms. In addition, alternative splicing regulation in exons 11, 12 and 13 yields another level of complexity, resulting in three different C-termini combinations within the F domain, constituting the highly variable C-terminal region typical of NR.

**Figure 1:**
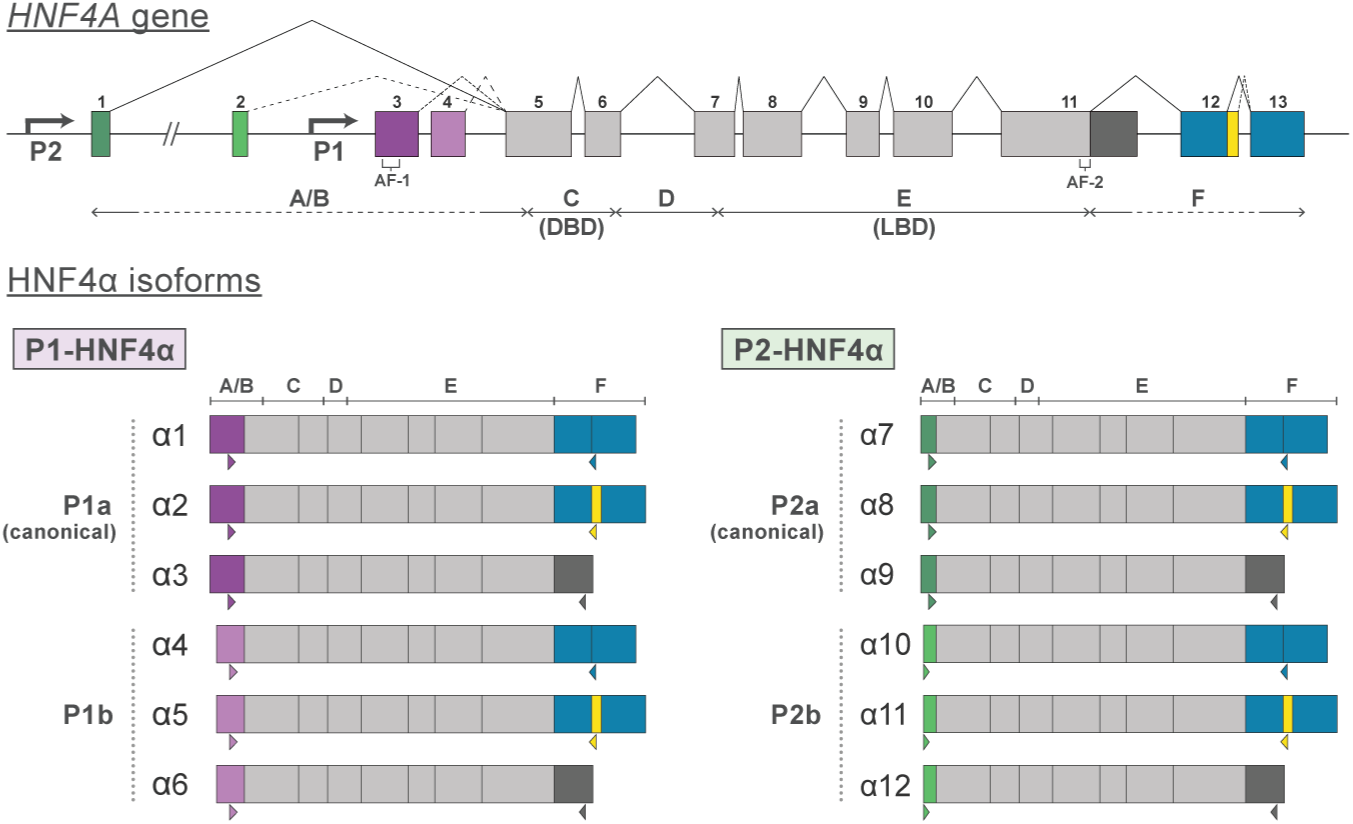
The HNF4A gene leads to the expression of 12 isoforms, through the use of alternative promoters and alternative splicing. The proximal P1 promoter can produce two distinct N-terminal ends through either exon 3 (P1a) or exon 4 (P1b), and the same number of isoforms from the distal P2 promoter are possible from either exon 1 (P2a) or exon 2 (P2b). Furthermore, alternative splicing at the C-terminal end can produce an additional three isoforms for each of these subgroups, resulting in twelve possible isoforms. The P1 and P2 promoters of the *HNF4A* gene are separated by approximately 46,000 bp.

Different numbering and identifiers are currently used when referring to the HNF4α isoforms (Supplementary Table 1), and several studies on HNF4α either do not precisely indicate which isoform was used, do not provide the cDNA sequence, or simply refer to a previous study. This has led to contradictory functions being attributed to HNF4α as a consequence of studying different isoforms (Babeu et al., 2018; Chellappa et al., 2016; Vuong et al., 2015). In order to simplify the nomenclature of the isoforms, we now propose to follow the P1 and P2 classification of isoforms, first by sorting the isoforms from the N-termini, and then by the different C-termini as also suggested by (Ko et al., 2019). This leads to four main classes of isoforms (P1a, P1b, P2a, P2b) (Figure 1), and maintains the order of the different C-termini in order to remain consistent with the nomenclature used in most previous studies. The isoforms a (P1a and P2a) represents the canonical HNF4α isoforms, whereas the isoforms b (P1b and P2b) are isoforms that have been less studied, or that have not been detected. In order to be consistent with the order of the C-terminal between the different isoforms, we also propose to inverse the nomemclature of the α4 and α5 isoforms (Drewes et al., 1996).

### Expression of HNF4α isoforms in different tissues

While multiple HNF4α isoforms have been considered in several studies, the expression of some of these isoforms has only been limited to a single study (α5 (Drewes et al., 1996), α10, α11 and α12 (Huang et al., 2009)), with α4 and α6 isoforms remaining putative, their expression having never been demonstrated. The expression of the different classes of HNF4α isoforms is tightly regulated depending to the spatial context and stage of development (Chen et al., 1994). Immunohistochemical analysis of several human tissues reveals a variable distribution of P1 and P2 isoform expression (Tanaka et al., 2006). The intestine is the only adult organ expressing both the P1 and P2 isoforms. The expression levels of these isoform classes have been modulated in the colon, notably by RNA interference in a colorectal cancer line or by exon exchange in the mouse, in order to analyze the functional differences between P1 and P2. These studies revealed significant disparities, with P1 isoforms being involved in the regulation of cell differentiation and metabolism, whereas P2 isoforms were associated with cell proliferation and cancer progression (Babeu et al., 2018; Chellappa et al., 2016).

To validate the existence of these 12 isoforms, different tissues of the human gastrointestinal tract were selected in the present study. A human cDNA library from healthy individuals was used to study isoform expression by semi-quantitative PCR (Figure 2), and the PCR products were sequenced to confirm the exact nature of the isoform. Seven isoforms were identified in these tissues (Figure 2). The canonical isoforms of HNF4α (P1: α1, α2, and α3; P2: α7, α8, and α9) were detected in the liver, stomach as well as all segments of the small intestine and colon. No isoforms were found in the esophagus, a tissue previously reported not to express HNF4α (Tanaka et al., 2006). Although the above approach was non-quantitative, a predominance of P1 isoforms in the liver was observed, consistent with the literature (Torres-Padilla et al., 2001). HNF4α5 was the only non-canonical isoform identified in this experiment, and found exclusively in the liver.

**Figure 2:**
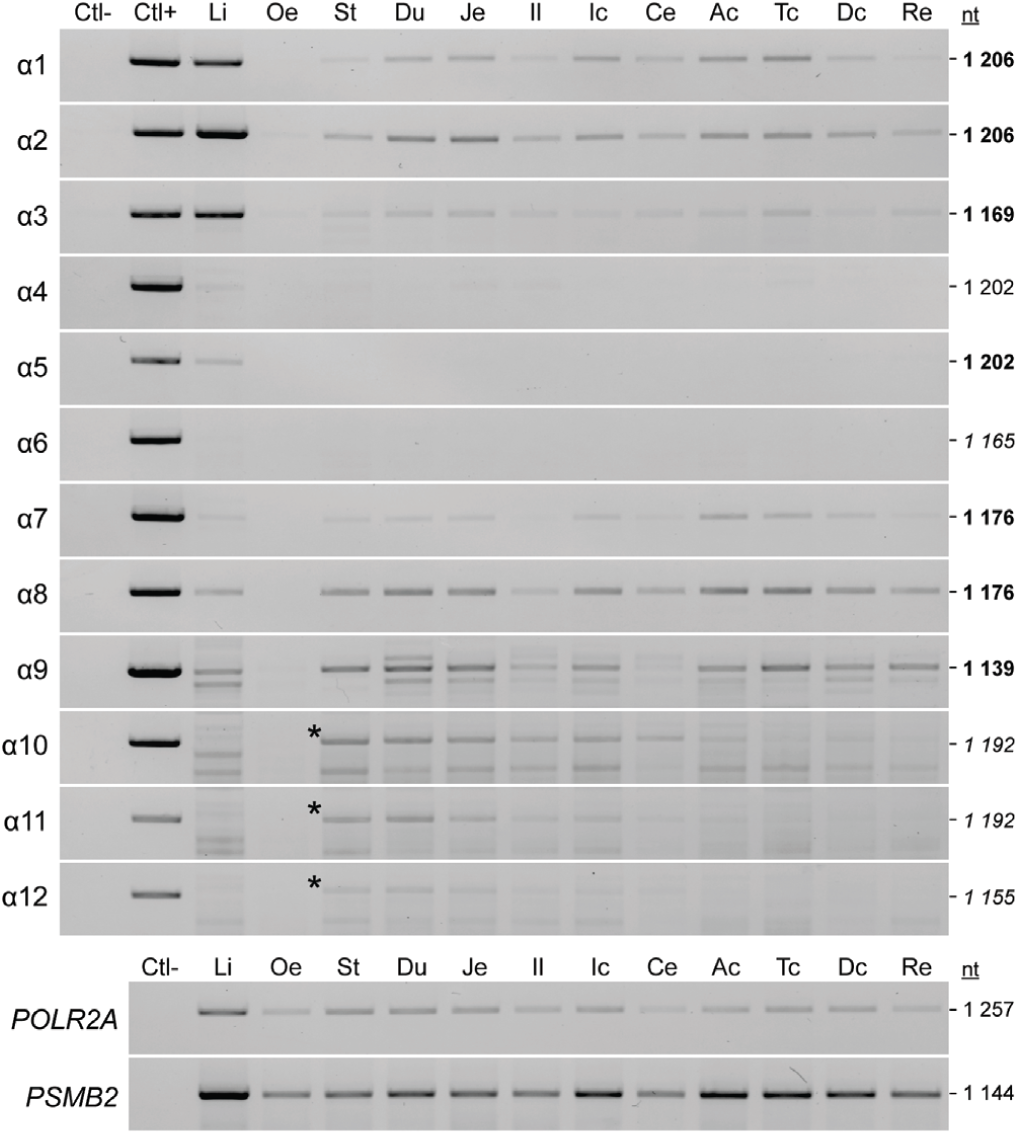
HNF4α isoforms are expressed at varying levels in different tissues of the human gastrointestinal tract. Expression of isoforms was assessed by semi-quantitative PCR in the liver, esophagus, stomach, and in various segments of the small intestine and colon. The α4, α6, α10, α11 and α12 isoforms were not detected in any of these tissues. The POLR2A and PSMB2 genes were used as reference genes.

Given that the expression of P1 and P2 HNF4α isoforms is known to be modulated in various cancer types (Tanaka et al., 2006), various human cancer lines were hence selected in an attempt to investigate the expression of additional isoforms. Two cell lines of pancreatic origin, AsPC-1 and Capan-2, were also included into this assay given that the pancreas is a well-known organ for expressing HNF4α. Expression of the 12 isoforms was measured in these cell lines by semi-quantitative RT-PCR (Supplementary Figure 1), and confirmed by sequencing of the PCR products. In the context of the cell lines selected for this assay, HNF4α1, α2, α3, α7, α8, and α9 were detected according to a pattern similar to the expression in the corresponding healthy human tissues.

### Generation of stable cell lines expressing inducible HNF4α with GFP and BioID2

Colorectal cancer cell lines that show endogenous expression of HNF4α unequivocally express several isoforms in a concomitant manner (Supplementary Figure 1). In order to study the specific functions of the different HNF4α isoforms, the latter had to be expressed in a cell line lacking endogenous expression of HNF4α. The HCT 116 cell line was ultimately selected for this purpose, since the *HNF4A* gene is mutated in this line, leading to loss of its expression (Barretina et al., 2012) (Supplementary Figure 1). The Flp-In T-REx system was used to generate stable lines in HCT 116 cells for each of the HNF4α isoforms, with either a GFP or a BioID2-3myc fusion protein at their C-terminal end (Figure 3A). This system allows integration of a gene at a single and known location within the genome, under the control of a doxycycline-inducible CMV promoter. In addition, this strategy supports gene expression levels that are much closer to an endogenous expression level comparatively to other transfection approaches, with these levels remaining relatively similar between the different stable cell lines.

**Figure 3:**
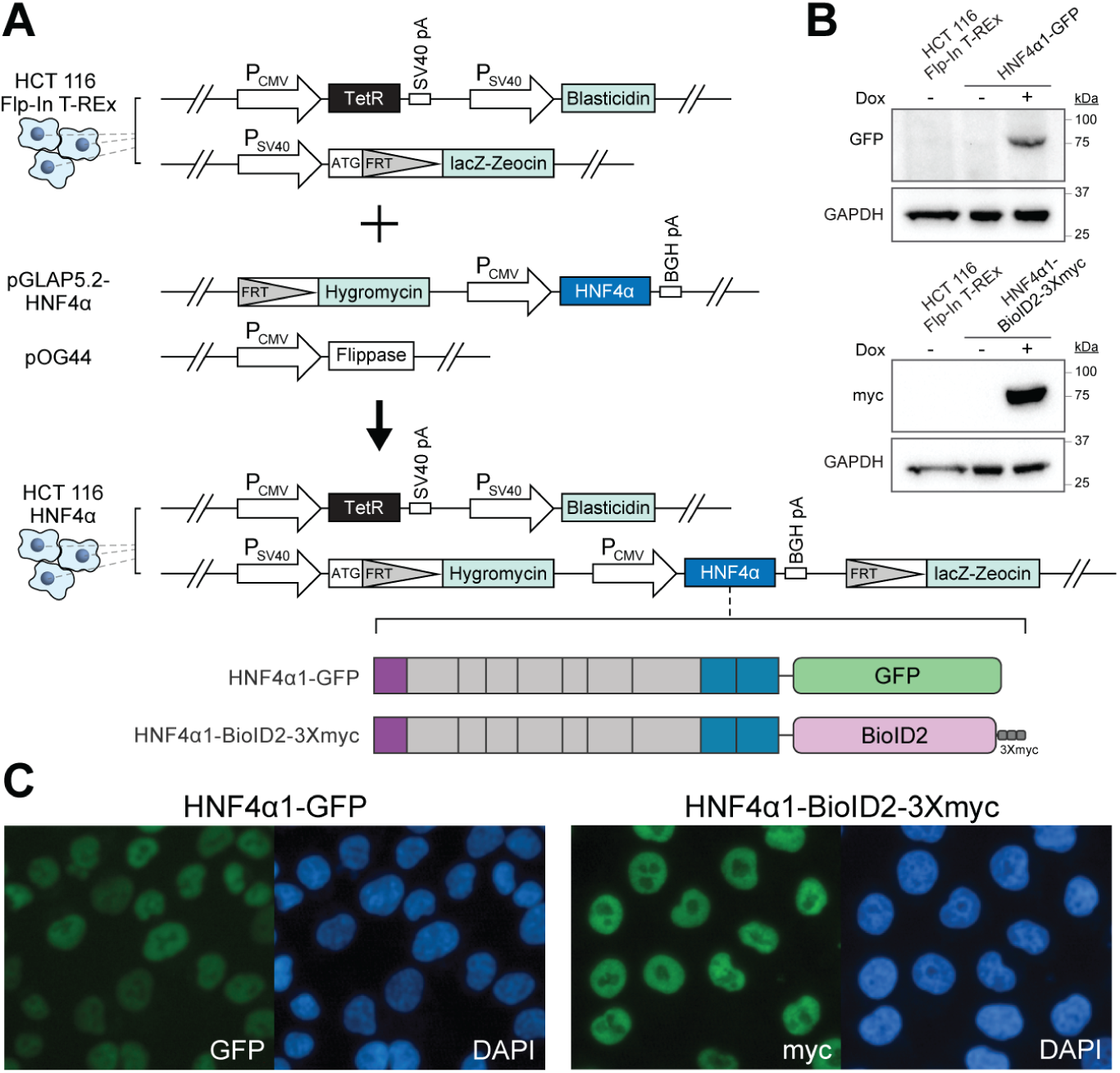
Generation of stable cell lines expressing inducible HNF4α with GFP and BioID2. **A)** The Flp-In T-REx system was used to generate stable lines in HCT 116 cells for each of the HNF4α isoforms, with either a GFP or a BioID2-3myc fusion protein at its C-terminal. The constructs of the 12 HNF4α isoforms fused with the GFP or BioID2-3myc protein labels were expressed following the induction of the stable lines in the HCT 116 cells. **B)** Immunoblots against the GFP and myc protein labels were performed to detect the expression of the different fusion proteins following induction with doxycycline of the cell lines stably expressing HNF4α1. The expression levels of the GAPDH protein in these total protein extracts were used as a reference. **C)** Nuclear localization of HNF4α1 was confirmed by immunofluorescence to detect the GFP tag (left) or the myc tag (right).

The inducible expression of HNF4α was confirmed by both immunoblotting (Figure 3B) and immunofluorescence microscopy when comparing cells incubated or not with doxycycline (Figure 3C). A similar nuclear localization was observed for all 12 isoforms, whether HNF4α was fused to a GFP (Supplementary Figure 2) or to BioID2-3myc protein tag (Supplementary Figure 3). Given the single integration site, the overall expression of HNF4α was expected to be relatively similar. However, differences in protein expression were observed when comparing immunoblot analysis of whole cell lysates (Supplementary Figure 4). This difference was most likely due to post-translational stability, since no correlative pattern of transcript expression was observed when measured by qPCR (Supplementary Figure 5).

### HNF4a isoforms have different DNA binding capacities

Following confirmation of the expression and localization of the 12 HNF4α isoforms, a first functional validation was performed by measuring their ability to bind the known consensus DNA binding sequence DR1 (Fang et al., 2012; Jiang and Sladek, 1997). Electrophoretic mobility shift assays (EMSA) were carried out to determine whether the 12 isoforms could bind this consensus DNA sequence (Figure 4A). Isoform binding to a probe containing the HNF4α DR1 response element resulted in a major complex shift for all isoforms except the α4, α5 and α6 isoforms (Figure 4A). Inclusion of an antibody against GFP in the binding reaction was able to supershift this complex demonstrating that the latter indeed consisted of the HNF4α isoforms fused to GFP. A probe featuring a mutated DR1 sequence was used to demonstrate the specificity of interaction of the isoforms with the consensus DR1 sequence (Figure 4A). This test demonstrates for the first time that the α4, α5 and α6 isoforms are unable to bind DNA in the same manner as other HNF4α isoforms, despite the fact that they share the same DNA binding domain. These results therefore suggest that the A/B domain from exon 4 of these isoforms can negatively regulate their DNA binding capacity, since this domain is the only common sequence for α4, α5 and α6 that is absent from the other nine HNF4α isoforms.

**Figure 4:**
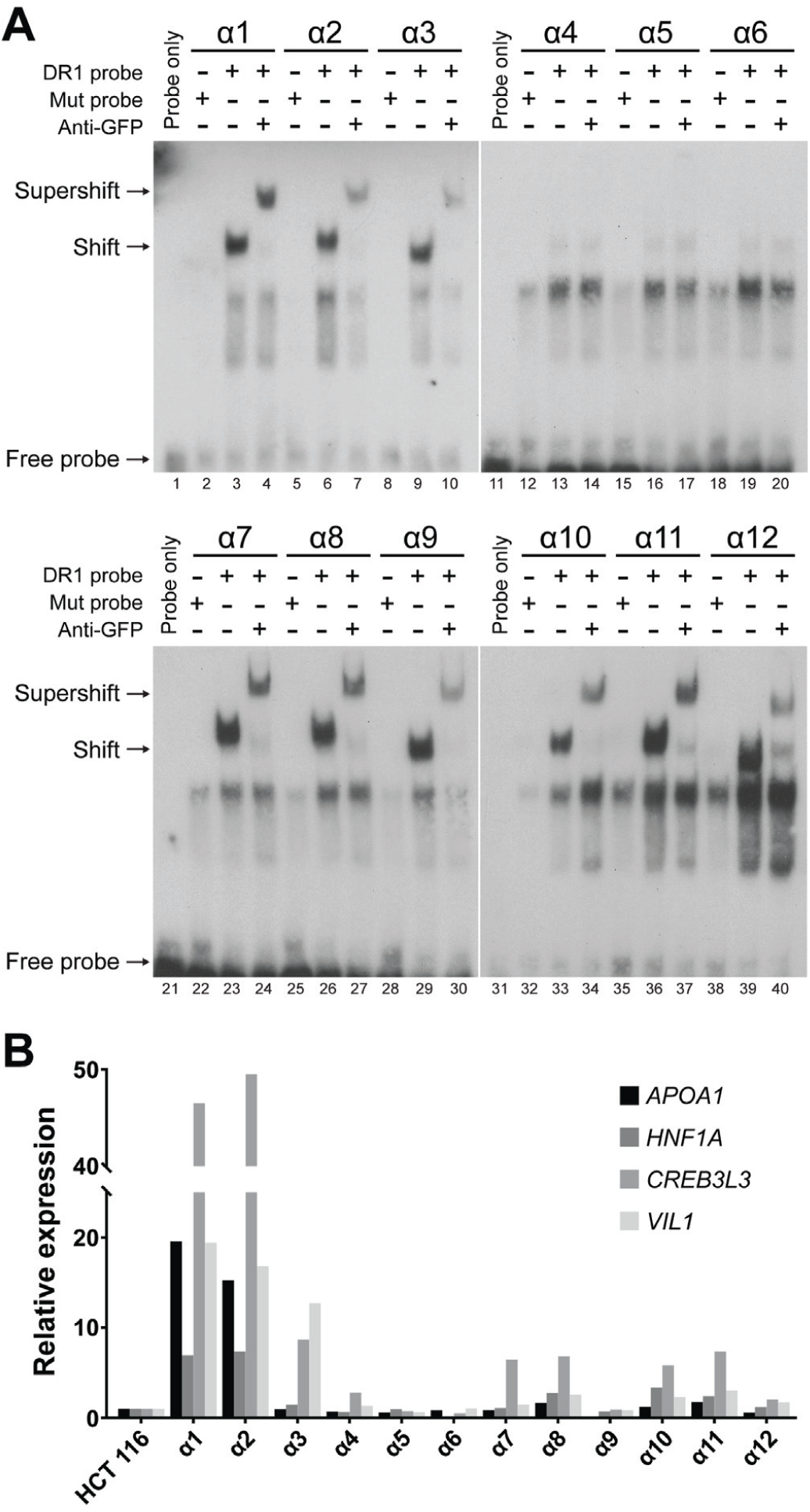
HNF4α isoforms do not all recognize the DR1 consensus response element of the nuclear receptor. A) 293T cells were transfected for 24 hours with the different HNF4α (1-12)-GFP constructs. Nuclear extracts from these cells were used to perform EMSA. A 5’-biotinylated DNA probe containing the normal or mutated DR1 consensus sequence of 5’-biotinylated HNF4α was incubated with each cell extract and the DNA-protein complexes were resolved by non-denaturing electrophoresis. Supershifts were performed using an antibody against GFP to confirm the presence of GFP-HNF4α in the observed complexes. The biotinylated probes were revealed with streptavidin-HRP. B) Known target genes of HNF4α were measured by quantitative RT-PCR reactions (qPCR) performed on samples from stable HCT 116 HNF4α (1-12)-GFP lines that were induced for 48 hours in the presence of 2.5 μg/ml doxycycline

HNF4α is known to transactivate the expression of a multitude of genes, the majority of which are involved in cell differentiation, metabolism and transport of nutrients. The expression levels of four selected genes known to be upregulated by HNF4α (*APOA1, CREB3L3, HNF1A* and *VIL1*) and involved in several of these functions were measured by qPCR following the induction of expression of each isoform in stable HCT 116 lines (HNF4α (1-12)-GFP) (Figure 4B). Significant differences in the transactivation of these genes were observed between isoforms. Although the isoform transactivation profiles were similar from one gene to another, there were a few notable exceptions. The α1 and α2 isoforms (canonical P1) were the strongest transcription inducers of the four genes tested, whereas the α3 isoform positively regulated *CREB3L3* and *VIL1* at lower levels, but not *APOA1* and *HNF1A* (Figure 4B), thus demonstrating specificity in the regulated genes between the different isoforms. The α4, α5 and α6 isoforms showed no effect on the genes tested, suggesting that the absence of binding to the DR1 element by these isoforms has a major impact on their transactivation activity (Figure 4A). The canonical (α7, α8) and non-canonical (α10, α11) P2 isoforms exhibited a much lower transactivation capacity than the α1 and α2 isoforms. Of particular note, for all analyzed genes, the variants of each subgroup sharing a shorter F domain (α3, α6, α9 and α12) all showed a strongly reduced or non-existent transactivation capacity for these genes. Importantly, while the isoforms exhibit differences in HNF4α protein expression (as demonstrated in Supplementary Figure 5), the results presented in Figure 4B show that these variations in protein expression are not necessarily consistent with the transcriptional levels of target genes.

### Transcriptomics analysis of HNF4a isoforms

In order to further study the transcriptional functions of the 12 HNF4α isoforms, the transcriptome of HCT 116 HNF4α (1-12) -GFP lines were analyzed by RNAseq. To achieve this, RNA from these lines were sequenced in triplicate, and compared to the parental cell line lacking HNF4α expression. The readings were aligned to the human transcriptome constructed from the annotations of the RefSeq database (Supplementary Table 1). The differential expression of transcripts and genes was obtained by comparing the specific data for each isoform to the transcriptome of the control condition (HCT 116 Flp-In T-REx).

Principal component analysis (PCA) of the datasets was performed to determine the variance found between the transcripts quantified for all isoforms as well as to validate the reproducibility of the triplicates for each experiment. This analysis confirmed the propinquity of each triplicate, with the exception of the first replica for the α3 isoform (symbolized by A3N1) (Figure 5A). The divergence of this replica is likely explained by a weaker expression of the HNF4α3 isoform in the sample (405 counts against 832 and 771 in the other two replicas). The subsequent analyses were therefore carried out by excluding this sample, and using the average value of the counts between the different biological replicas. The control condition is annotated as A0, and is located on the left and center of the PC1 and PC2 components respectively. From these analyses, the α1 and α2 isoforms caused the most variance when compared to control, while α3 exhibited a reduced effect. In contrast, the α4, α5 and α6 isoforms caused very little divergence. The P2 isoforms displayed an intermediate effect with respect to these two subgroups of P1 isoforms. Of note, the isoforms generally appeared to cluster strongly according to their A / B domain, with the exception of a larger divergence observed between the α10, α11 and α12 isoforms, indicating that the majority of the functional differences between the HNF4α isoforms were associated with this domain rather than with the F domain. Notwithstanding the latter, a shorter F domain effect contained in the α3, α6, α9, α12 isoforms resulted in closer proximity to the negative control than to the other two isoforms containing the same A / B domain.

**Figure 5:**
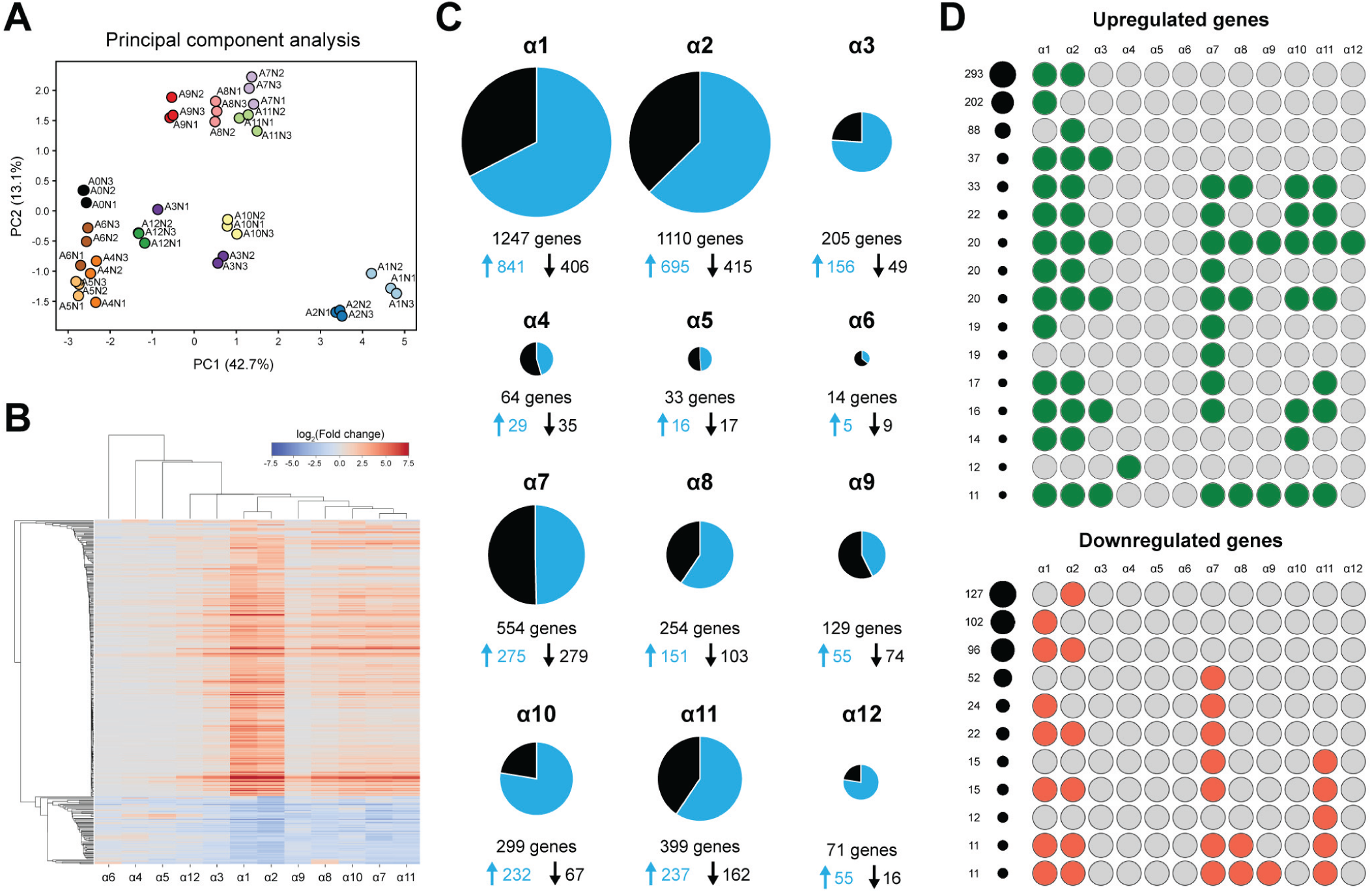
Transcriptomics analysis of HNF4a isoforms by RNAseq. **A)** PCA analysis of the RNAseq data from GFP cell lines expressing each of the 12 HNF4α isoforms (A1-A12) and the control condition (A0). The biological triplicates for each sample are represented by N1/N2/N3 following the isoform number. The two main components PC1 and PC2 were used for two-dimensional visualization of the analysis. **B)** Hierarchical analysis of the cell lines expressing the different HNF4α (1-12)-GFP were compared based on their RNA expression profiles using the log_2_ fold change. **C)** Representation of the number of transcripts showing significant up and down regulation for each of the HNF4α isoforms. A minimum absolute modulation threshold of 2 combined with an adjusted p-value threshold ≤ 0.001 was used to determine significance. The size of the circles is proportional to the total number of genes modulated for each isoform. The blue color represents the proportion of up-modulated genes relative to the control condition, while the black-colored portion represents the negatively modulated genes. **D)** Graphical representation with the number of regulated genes common to the different isoforms or unique to specific isoforms, with green indicating upregulation and red indicating downregulation. Only groups with more than 10 genes are shown.

Gene quantification analysis revealed that the annotation contained ∼45000 genes, with ∼18000 detected in at least one sample (using a minimal quantification of 1 TPM). To further analyze the observed variance, only those genes positively or negatively modulated by at least two-fold compared to the control, and with a minimum Benjamini-corrected p-value of 0.001, were considered. The results are viewed globally in Figure 5C as circles whose respective sizes are proportional to the number of significantly modulated genes. Positive and negative modulation levels are also presented as volcano plot representations of the sorted results in order to visualize the distribution of the genes found across the different data sets (Supplementary Figure 6). The most striking difference between the transcriptome profiles induced by the 12 HNF4alpha isoforms was the total number of modulated genes (Figure 5C). The α1 and α2 isoforms were by far the isoforms with the greatest impact on the transcriptome in this setting. The α4, α5 and α6 isoforms conversely displayed an extremely weak effect on gene expression, consistent with the absence of DNA binding to the DR1 consensus sequence. This observation also suggests that they do not bind to a different response element with functional impact on the transcriptome. An intermediate effect was noted for the canonical and non-canonical P2 isoforms. In general, HNF4α was found to be more involved in the activation of gene expression than in their repression. The genes modulated by the α1 and α2 isoforms, which are the most important regulators in this setting, were mostly modulated upward (67% and 63%, respectively). Certain isoforms such as α9 were mainly responsible for repression of transcription (57%), while others such as α3 had an even stronger activating effect (76%), but with a much less overall number of genes regulated (Figure 5C). Comparison of the overlap of upregulated or downregulated genes between the different isoforms showed the largest group being regulated by α1 and α2 isoforms (293 genes), although a large number of genes were also specific to α1 (202 genes) (Figure 5D). Of note, most of the genes regulated by the P2 isoforms were common to all isoforms in this group and were also regulated by at least one P1 isoform, whereas the few genes regulated by the α4-5-6 isoforms were completely different from the remaining isoforms (Figure 5D). Overall, these data demonstrate major differences in transcriptional function between the 12 isoforms of HNF4α.

### HNF4α isoforms have distinct interaction networks, mostly comprised of transcription factors and transcriptional coregulators

The functional activity of HNF4α is regulated by two characteristics related to its structure. Its DBD mediates the recognition of specific regulatory sequences at its target genes, while different transactivation domains promote the interaction with multiple transcriptional coregulators. Since the 12 isoforms have the same DBD, the differences in the number of modulated genes observed between the HNF4α isoforms likely arise from a variable interaction of the isoforms with transcriptional coregulators via the transactivating A/B or F domains. In order to determine the specific protein-protein interaction networks involving each of the HNF4α isoforms, a BioID approach coupled to SILAC (Stable isotope labeling with amino acids in cell culture)-based quantitative mass spectrometry was used (Figure 6A) (Varnaite and MacNeill, 2016). Stable HCT116 cell lines were generated from constructs expressing each of the 12 HNF4α isoforms in fusion with BioID2 (Kim et al., 2016). Three conditions were thus compared using SILAC, namely the control cell line in light medium (R0K0), a cell line expressing a P1 isoform in the intermediate SILAC medium (R6K4) and a line expressing a P2 isoform in heavy medium (R10K8) (Figure 6A). The biotin-labeled protein extracts from these three conditions were then mixed in a ratio of 1: 1: 1. The biotinylated proteins were digested with trypsin, and the resulting peptides analyzed by mass spectrometry. Quantification of the identified proteins for each isoform was performed by measuring the enrichment in comparison to the control condition (Figure 6A).

**Figure 6:**
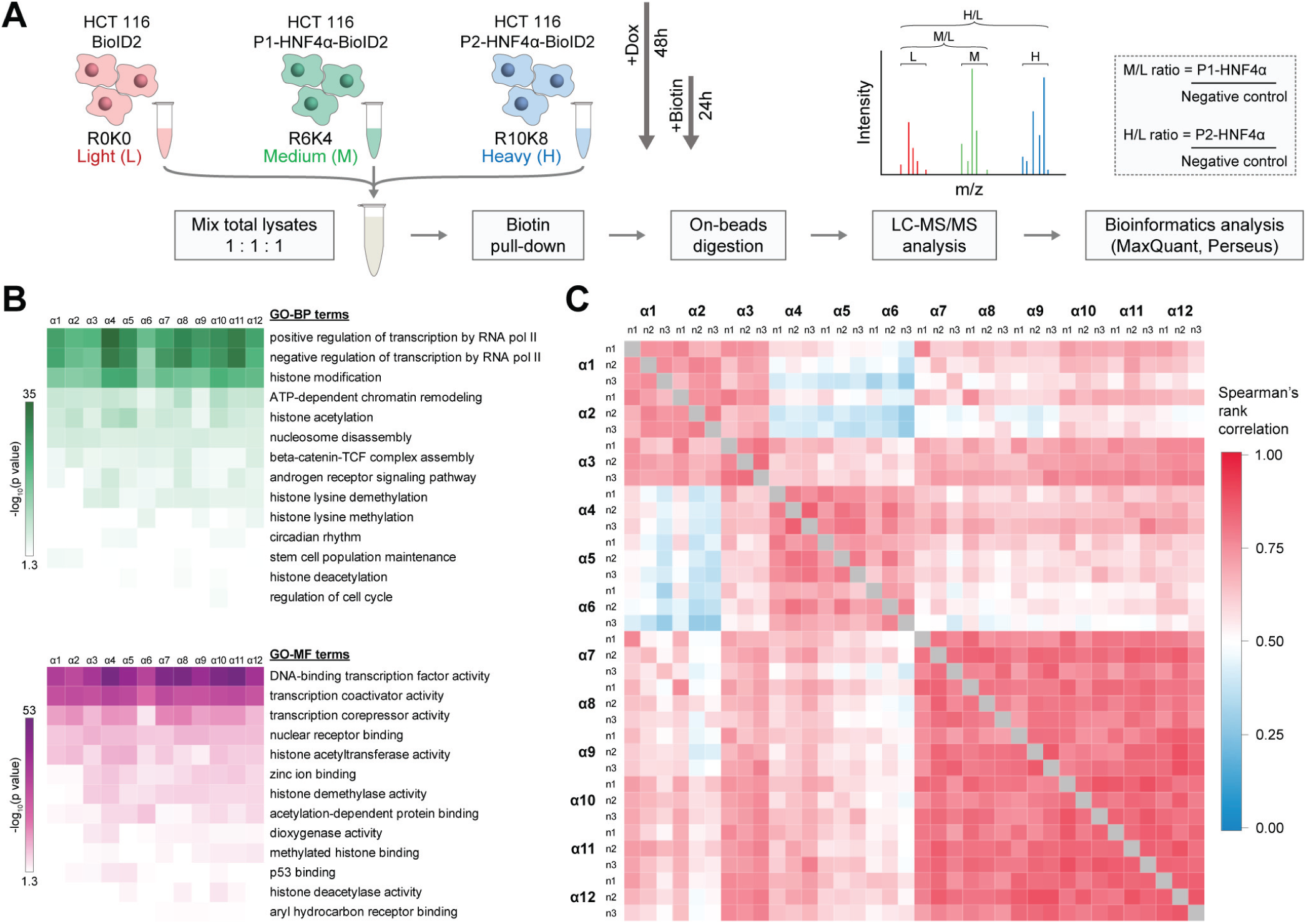
Identification of protein interactions of HNF4α isoforms using the BioID approach coupled with quantitative mass spectrometry. **A)** The HCT 116 HNF4α (1-12) -BioID2-3myc and the HCT 116 control line BioID2-3myc-empty cell lines were cultured in SILAC medium, induced for 48 hours with doxycycline and incubated for 24 hours with biotin. The total extracts from these cells were combined equally. The biotinylated proteins were precipitated with streptavidin-coupled beads and then trypsin-digested on beads. The peptides were analyzed by LC-MS/MS, before quantifying the results using MaxQuant. The experiments were performed in triplicates **B)** The identified interacting proteins were analyzed for gene ontology enrichment for biological processes (GO-BP, green) and molecular functions (GO-MF, purple) annotations for the proteins identified by each of the HNF4α isoforms. The Panther 13.1 tool was used for the above analyses while the Morpheus software was used to visualize the results as a heatmap. A minimum enrichment ratio threshold of 2 was used. The negative value of the logarithm in base 10 of the p-value is represented according to a color scale for each annotation. **C)** Diagram showing the enrichment ratios according to the experimental conditions used for culture in SILAC medium. Heatmap visualization comparing the association between the protein enrichment ratios identified for each isoform according to a Spearman correlation. The data are presented according to the biological triplicates carried out for each isoform.

Over a thousand proteins were identified for each isoform. Enrichment ratios obtained for each quantified protein were compared between isoforms and their triplicates via a Spearman correlation. These correlations were then plotted against a color scale for easier visualization (Figure 6B). This initial analysis validated the reproducibility of the obtained results, yielding a very strong correlation between the triplicates (0.72 to 0.95). In addition, this representation highlighted certain differences between the proteomes associated with each isoform. The lists of potential interactants for the α4, α5 and α6 isoforms were the most dissimilar compared to the other isoforms. Overall, it appeared that most of the identified proteins were similarly enriched for the 12 isoforms. Following this validation, the remaining analyses were carried out based on the average enrichment ratios obtained through the triplicates. Given the differential isoform expressions observed (Supplementary Figure 4), the enrichment ratios were normalized relative to the median.

In order to determine the type of proteins enriched by the BioID approach, enrichment analyses of GO annotations were carried out using the Panther 13.1 software. A minimum enrichment threshold of 2 was used for these assays, reducing the number of proteins to approximately 200 interactants for each isoform. Annotations of biological processes (GO-BP) regulated by these proteins showed an enrichment in the regulation of transcription, notably through histone modification and chromatin remodeling (Figure 6C). As expected, the proteins associated with these enriched biological processes were annotated with molecular functions (GO-MF) for transcription factors and transcriptional coregulators (Figure 6C). There were, however, no significant differences between the proteomes of the 12 isoforms that could be identified by this type of analysis, demonstrating that all of these isoforms interact with proteins with known functions in transcription.

### Identification of isoform-specific interaction partners of HNF4α

In order to determine whether specific interactors could be identified for each isoform, the respective effects of the A / B domain and F domain were decoupled. Four different A / B domains and three F domains are found among all isoforms. By pooling the identified proteins for all of the isoforms having one of these domains in common, this analysis enabled to provide clues as to the identification of proteins that can interact specifically with certain HNF4α domains generated by alternative splicing.

The Venn diagrams presented in Figure 7 illustrate the results of this comparative analysis, which used a minimum enrichment threshold of 2. Overall, most of the identified proteins were once again enriched equally between the 12 HNF4α isoforms, which included 69 proteins that were enriched by a 2-fold ratio for all isoforms, in all triplicates. Among these, there were several coregulators and transcription factors known to interact with HNF4α, such as CBP (Dell and Hadzopoulou-Cladaras, 1999), NCOA-1 and NCOA-2 (Martinez-Jimenez et al., 2006), NCOR2 (Ruse et al., 2002) and FOSL1 (FRA-1, (Vuong et al., 2015)). This analysis, however, highlighted certain proteins that appeared to interact specifically with subgroups of isoforms that had a common A/B domain or F domain. Most of these proteins have never been shown to specifically interact with HNF4α, although their functions in transcriptional regulation strongly support this possibility. These include members of ATP-dependent chromatin remodeling complexes (BRD9 - SWI / SNF, VPS72 - NuA4, SRCAP), enzymes associated with histone modification (SIRT1, SIN3B, KDM2A, KANSL1), and members of the Mediator and PIC complexes (MED26, MED27, TAF4B). Comparison of proteins interacting with the different HNF4α F domains also demonstrated specific interactions with the three different C-terminal domains (Figure 7B), including the transcriptional corepressors IRF-2BP1 and IRF-2BP2.

**Figure 7:**
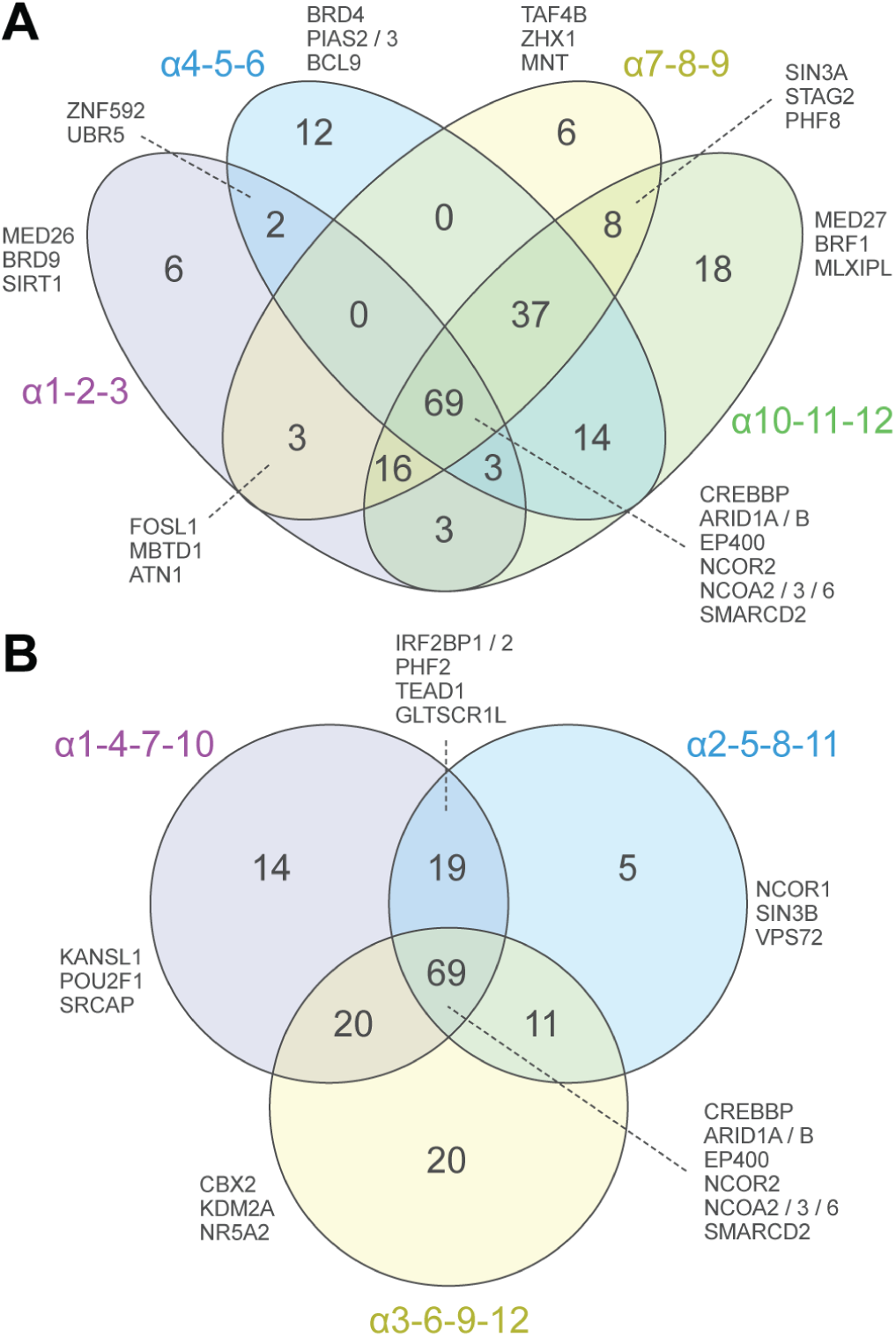
Several proteins specifically interact with isoforms that have a common A / B or F domain. Venn diagrams comparing specifically identified proteins in certain subgroups of HNF4α isoforms. A minimum enrichment ratio threshold of 2 was used. **(A)** Comparison of isoform-specific interactants sharing the same A/B domain. **(B)** Comparison of isoform-specific interactants sharing the same F domain.

To confirm some of these interactions, co-transfections between these newly identified FLAG-tagged interactants and GFP-tagged HNF4α isoforms showing the highest SILAC enrichment ratio were performed. Following immunoprecipitation against GFP, immunoblotting against FLAG led to the validation of the interactions involving five proteins, namely GATAD2B (α11), HNF4γ (α4), IRF2BP2 (α2), MTA1 (α11) and ZNF629 (α11) (Supplementary Figure 7). The control condition, which only expressed the FLAG construct, confirmed the specificity of interaction between HNF4α and these proteins. Thus, the identification of interactions by BioID coupled to SILAC quantification provided novel information with regard to the transcription factor interaction networks of HNF4α.

## DISCUSSION

*HNF4A* has been reported to produce several isoforms through the use of two different promoters and various alternative splicing events (Babeu and Boudreau, 2014). Two major groups of isoforms were classified as P1- and P2-HNF4α, according to the usage of their specific promoter. Given the presence of four different N-terminal regions, we have proposed a novel nomenclature based on these differences. Although we were successful in detecting seven of the twelve possible *HNF4A* isoform transcripts among specific gastrointestinal tissues and cell lines, we cannot exclude that the remaining isoforms are specifically produced in other tissues or in a timely manner during development. Despite several of these transcripts being shown to produce proteins (Tanaka et al., 2006), the repetitive and combinatorial nature of A/B and F domains in certain isoforms makes it impossible to detect single protein isoforms. However, the generation herein of a cellular system with controlled expression for each recombinant single isoform led to the demonstration that each isoform produces protein with possible differing post-translational stability as previously reported for subclasses of P1 isoforms (Chellappa et al., 2016). Since functional differences in the cell transcriptome were identified among each HNF4α isoform, our study supports the importance for the design of thorough analyses to measure the biological impact of each single isoform in a specific cellular and developmental context.

Although all HNF4α isoforms share the same DBD, our data support that P1B isoforms (α4, α5 and α6) do not functionally interact with DNA or influence gene expression when expressed as single isoforms producing homodimers. Our BioID approach coupled to quantitative mass spectrometry identified several transcriptional co-regulators interacting with these isoforms similarly to other HNF4α isoforms. Since the common difference between these three isoforms resides in their A/B domain, we speculate that this region could interfere with DNA binding and down-modulate the transcriptional influence of P1B isoforms on gene transcription. Of note, similar observations were made for two RAR-related orphan alpha (RORα) nuclear receptor isoforms where RORα1 was shown to strongly activate transcription while RORα2 interaction with RORE elements was more restricted with the result of weaker transcriptional activity (Giguere et al., 1994). Similarly to HNF4α-P1A and B isoforms, RORα2 and 1 differ in their A/B domain. Whether the HNF4α-P1B subclass of isoforms act as dominant negative regulators for other HNF4α isoforms in a given cellular condition will require further investigation.

HNF4α was initially considered to be an exclusive homodimer, (Jiang et al., 1995). However, recent evidence demonstrate that HNF4α can also form heterodimers, with distinct gene targets from the corresponding homodimers (Ko et al., 2019), and several tissues and cell lines express more than one isoform (Figure 2, Supplementary Figure 1 and (Ko et al., 2019)). The overexpression of different homodimers and heterodimers appears to regulate differentially a subset of previously reported genes, as well as inflammatory genes, underlining the importance of further studies that will tackle the complete transcriptome regulated by different combinations of HNF4α isoforms.

The BioID quantitative mass spectrometry performed for each HNF4α isoform identified a prominent number of common proteins that constitute ATP-dependent chromatin remodeling complexes such as the BRG1/BRM-associated factor (BAF) and the nucleosome remodeling and deacetylase (NuRD) complex (Supplementary Figure 8). These observations suggest that all HNF4α isoforms have the potential to act on chromatin structure via these specific interactions. Our results also suggest that the A/B domain is the major determinant of the differential transcriptional function of the different isoforms. However, we also observed differences in protein interactions such as IRF-2BP2. This protein was initially shown to be a corepressor of the IRF-2 transcription factor, and more recently for other transcription factors such as p53 and NFAT (Carneiro et al., 2011; Childs and Goodbourn, 2003; Koeppel et al., 2009). The BioID approach used herein show that isoforms that have the short form of the F repressor domain do not interact with this coregulator. Taking into account the differential interaction profile of IRF-2BP2 with the HNF4α isoforms, we have identified transcriptome variations that can be explained by these preferential interactions. Accordingly, our results show that the proportion of down-modulated genes was similar between the α1 and α2 isoforms (33% and 37%, respectively). This proportion fell to 24% at the level of the α3 isoform, with a total of approximately 8 times less negatively modulated genes. Since the IRF-2BP2 corepressor was shown to specifically interact with α1 and α2 isoforms relative to α3, it is possible that IRF-2BP2 may be responsible for part of this variation in the negative transcriptional regulatory function of these isoforms.

In conclusion, the present study provides an exhaustive analysis of the transcriptional co-interacting complexes and transcriptomic impact of each single HNF4α isoforms in a specific cellular setting. Given that specificity of biological functions was previously attributed to HNF4α-P1 and -P2 subclasses (Babeu et al., 2018; Chellappa et al., 2016), this work provides a strong rationale to further detail the exact nature of contributing HNF4α isoforms in given biological systems and to design innovative strategies in exploring the specific biological functions that may be coincidental to the expression of these specific isoforms in the biological field. Similar analyses could be designed for the study of other human NR isoforms that result from similar alternative promoter and splicing events.

## EXPERIMENTAL MODEL AND SUBJECT DETAILS

All cell lines were obtained from the ATCC. HCT 116 (human colorectal cancer (hCRC)), Caco-2/15 (hCRC), Capan-2 (human pancreatic adenoma), HepG2 (human hepatocellular carcinoma) and 293T (transformed human embryonic kidney) cell lines were cultured in DMEM. AsPC-1 (human pancreatic adenocarcinoma), COLO 205 (hCRC) and DLD-1 (hCRC) cell lines were cultured in RPMI. The T84 (hCRC) cell line was cultured in DMEM/F-12. The LoVo (hCRC) cell line was cultured in F-12K. The HT-29 (hCRC) cell line was cultured in McCoy’s 5A. All cultured media were supplemented with 10% FBS and cell lines were grown in a humidified incubator at 37^0^C with 5% CO_2_. The Human Digestive System MTC Panel cDNA library was purchased from Clontech Laboratories (Mountain View, USA). The cDNA preparations found in this library are derived from the combination of cDNAs from several healthy Caucasian individuals between 18 and 61 years of age, and were provided at a concentration of 1.0 ng/μl following first-strand cDNA preparation for each tissue. A total of 12 tissues of the digestive system were included in the bank: liver, stomach, esophagus, duodenum, jejunum, ileum, ileum, cecum, ascending, transverse and descending colon, as well as the rectum.

## METHOD DETAILS

### Total RNA isolation and Reverse Transcription

Total RNA of HCT 116, Caco-2/15, T84, COLO 205, LoVo, DLD-1, HT-29, HepG2, AsPC-1, Capan-2 and HCT 116 HNF4α (1-12) -GFP cell lines was isolated using RNeasy RNA isolation kit (QIAGEN). The stable HCT 116 HNF4α (1-12) -GFP cell lines were induced for 48 hours with 2.5 μg/ml doxycycline (Clontech Laboratories, Mountain View, USA) prior to extraction of total RNAs. cDNA synthesis was performed with the SuperScript IV-RT reverse transcriptase enzyme (Thermo Fisher Scientific, Waltham, USA). Two μg of RNA were added in a total volume of 10 μl by supplementing with DEPC water. A mixture containing 2.4 μl of 0.5 μg/μl poly (dT) oligos (Amersham Biosciences, Little Chalfont, United Kingdom) and 0.8 μl of 25 mM dNTPs (Amersham Biosciences, Little Chalfont, United Kingdom) was added to the RNA, then heated for 5 minutes at 75°C and placed on ice for an additional 5 minutes. A 10 μl volume of RT reaction mixture, described in Table 4, was added to the RNA. The reaction was incubated for 1 hour at 50°C, before inactivating the SuperScript IV-RT by heating for 5 minutes at 95°C. The cDNA samples were subsequently stored at −20°C.

### Expression of isoforms in different human cancer cell lines

Oligonucleotides specific for each isoform and the reference genes were obtained from Integrated DNA Technologies (IDT, San Jose, USA) (Supplementary Table 6). The cDNA from HCT 116 HNF4α (1-12) -GFP lines was used as a positive control for each isoform. Expression levels of the HPRT and PUM1 genes were used as references, with an amplification of 26 cycles at a hybridization temperature of 60°C and an elongation time of 20 seconds.

### Expression of isoforms in the different tissues of the human digestive system

The expression of the 12 HNF4α isoforms in the different human gastrointestinal tract tissues was assessed by PCR. The primers specific to each HNF4α isoform (Supplementary Table 6) were used to assess the expression of the isoforms by PCR in each of the aforementioned listed tissues. The reagents used for the amplification were the same as described previously, the template DNA being in this instance a volume of 3.7 μl of cDNA. The PCR reactions were performed according to the PCR conditions detailed in the previous section, for a first round of 30 cycles. A 10 μl aliquot of the reaction was retrieved, and the remainder of the reaction was supplemented again at 20 μl at the initial reaction concentrations for a second round of PCR of 6 additional cycles. The expression levels of the POLR2A and PSMB2 genes were used as references, with an amplification of 26 cycles. A PCR product corresponding to each isoform was sequenced to ensure the specificity of the amplification. Plasmid pUC19 was digested with the SmaI restriction enzyme (New England Biolabs, Ipswich, USA) in CutSmart buffer (New England Biolabs, Ipswich, USA) for 1 hour at 25°C, and subsequently purified on gel agarose. Ligation between the PCR product and the digested pUC19 plasmid was carried out in a 20:1 ratio (insert: vector) with T4 DNA ligase (New England Biolabs, Ipswich, USA) for 3 hours at room temperature. Sequencing was performed via the Genome Sequencing and Genotyping Platform (Université Laval, Quebec, Canada).

### Cloning of HNF4α isoforms

The 12 HNF4α isoforms were cloned into the donor vector pENTR11 (Thermo Fisher Scientific, Waltham, USA) in three distinct steps consisting in initially cloning the sequence common to the 12 isoforms, inserting the different N-terminal termini (A/B domains) on each side, followed by the different C-termini (F domain). The Gateway system (Invitrogen, Carlsbad, USA) was subsequently used to obtain constructs allowing the expression of isoforms in fusion with the GFP and BioID2 protein labels.

#### Cloning of the common sequence in pENTR11

The common sequence to the 12 isoforms was amplified by PCR from the pLenti-HNF4α2-GFP plasmid. This plasmid contains the α2 isoform of HNF4α (NM_000457.3) synthesized by Feldan Inc. and cloned into pLenti6/V5 (Invitrogen). The amplified common sequence measures 1014 base pairs, covering 71-86% of the complete sequence of the different HNF4α isoforms, and contains the C, D and E domains of HNF4α. Oligonucleotides used for amplification (Supplementary Table 6) were obtained from Integrated DNA Technologies (IDT, San Jose, USA). These enabled the addition of SpeI and SfoI restriction sites within the common sequence, at the 5’ and 3’ ends, respectively, without changing the amino acid composition of the HNF4α protein. In parallel, these primers also allowed the upstream and downstream addition of the sequence of SalI and XhoI restriction sites, in consecutive order. The amplification reaction was performed with iProof High-Fidelity DNA Polymerase enzyme (Bio-Rad, Hercules, USA) according to the manufacturer’s recommendations. The reactions were performed in the T100 thermal cycler (Bio-Rad, Hercules, USA).

The PCR product was purified on agarose gel using the EZ-10 Spin Column DNA Gel Extraction Kit (Bio Basic, Markham, Canada). The purified PCR product and the pENTR11 plasmid were digested with SalI and XhoI restriction enzymes (New England Biolabs, Ipswich, USA) in NEBuffer 3.1 digest buffer (New England Biolabs, Ipswich, USA) for 2 hours at 37°C, then purified on agarose gel. The digested and purified pENTR11 plasmid was then dephosphorylated by the Antarctic Phosphatase enzyme (New England Biolabs, Ipswich, USA) for 30 minutes at 37°C, before ligation between the PCR product and pENTR11 in a 5: 1 ratio (insert: vector) with T4 DNA ligase (New England Biolabs, Ipswich, USA) for 2 hours at room temperature. The pENTR11-(common sequence of HNF4α) plasmid was sequenced via the Genome Sequencing and Genotyping Platform (Laval University, Quebec, Canada).

#### Inserting the N-Termini into pENTR11

The four different N-terminal ends of the HNF4α isoforms, surrounded by SalI (5’) and SpeI (3’) restriction sites, were obtained from Integrated DNA Technologies (IDT, San Jose, USA). The sequences corresponding to the N-terminal ends of the α1/2/3 and α4/5/6 isoform groups were synthesized directly in the form of double-stranded DNA (gBlocks Gene Fragments). The sequences corresponding to the N-terminal ends of the α7/8/9 and α10/11/12 isoform groups were obtained in the form of single-stranded oligonucleotides. In order to obtain a double-stranded sequence, the sense and antisense oligonucleotides were mixed at a concentration of 1 μM in a 5 mM NaCl buffer, 20 mM Tris pH 7.5. Hybridization of the oligonucleotides was performed by placing the reaction mixture in a heated dry bath set at 98°C for 2 minutes and subsequently allowing the reaction to return to room temperature after turning off the bath. The pENTR11-(common HNF4α sequence) plasmid as well as the four N-termini were digested with SalI and SpeI restriction enzymes (New England Biolabs, Ipswich, USA) in CutSmart buffer (New England Biolabs, Ipswich), USA) for 2 hours at 37°C, before being purified on agarose gel. A ligation reaction with a 10: 1 ratio (insert: vector) with T4 DNA ligase for 2 hours at room temperature was then performed to insert each N-terminus upstream of the common sequence of HNF4α already present in pENTR11. The four different plasmids thus obtained were sequenced via the Genome Sequencing and Genotyping Platform (Laval University, Quebec, Canada).

#### Inserting the C-Termini into pENTR11

The three different C-terminal ends of the HNF4α isoforms, lined with a 3’XhoI restriction site, were synthesized directly as double-stranded DNA (gBlocks Gene Fragments) (IDT, San Jose, USA). These three sequences as well as the four pENTR11 plasmids containing the different N-terminal ends in front of the common HNF4α sequence were digested by the restriction enzymes SfoI (only for the plasmids) and XhoI (New England Biolabs, Ipswich, USA) in the CutSmart buffer for 2 hours at 37°C, then gel purified. A ligation reaction as described previously was subsequently performed to insert each C-terminus downstream of the common HNF4α sequence into pENTR11. The pairing of three C-terminal ends at the four N-termini clones previously cloned hence generated 12 pENTR11-HNF4α plasmids (1-12), containing the complete sequence of each of the 12 HNF4α isoforms. These plasmids were sequenced via the Genome Sequencing and Genotyping Platform (Laval University, Quebec, Canada).

### Generation of pcDNA-DEST47, pgLAP5.2-GFP and pgLAP5.2-BioID2-3myc

The sequences of the 12 isoforms cloned into the pENTR11 vector were transferred into the pcDNA-DEST47 expression vectors (Thermo Fisher Scientific, Waltham, USA), pgLAP5.2 (a gift from Peter Jackson (Addgene plasmid #19706)) and pgLAP5.2-BioID2-3Xmyc by Gateway cloning, via an LR reaction according to the manufacturer’s instructions (Thermo Fisher Scientific, Waltham, USA). The pcDNA-DEST47-HNF4α (1-12), pgLAP5.2-HNF4α (1-12) and pgLAP5.2-HNF4α (1-12) -BioID2-3Xmyc plasmids were sequenced via the Genome Sequencing and Genotyping Platform (Laval University, Quebec, Canada). The control empty pGLAP5.2-BioID2-3Xmyc-plasmid was cloned in order to express the BioID2-3Xmyc protein label alone as a control for mass spectrometry experiments. To achieve the latter, the pGLAP5.2-BioID2-3Xmyc plasmid was amplified by PCR so as to remove an approximately 1.8 kb fragment containing the chloramphenicol resistance gene and the ccdB gene.

### Generation of inducible stable cell lines

The stable cell lines HCT 116 HNF4α (1-12) -GFP, HCT 116 HNF4α (1-12) BioID2-3Xmyc and HCT 116 BioID2-3Xmyc-control were generated using the Flp-In T-REx system (Thermo Fisher Scientific, Waltham, USA) using the pgLAP5.2-HNF4α (1-12), pgLAP5.2-HNF4α (1-12) - BioID2-3Xmyc and pGLAP5.2-BioID2-3Xmyc-empty plasmids, respectively.

Transfections for stable lines were performed with HCT 116 Flp-In T-Rex cells, upon reaching approximately 70% confluence, in a 60 mm Petri dish using Lipofectamine LTX (Thermo Fisher Scientific, Waltham, USA) as a transfection agent. A total of 500 ng of plasmid DNA was mixed with 4.5 μg of the Flp-Recombinase expression vector pOG44 (Thermo Fisher Scientific, Waltham, USA), 5 μl of Plus Reagent (Thermo Fisher Scientific, Waltham, USA) and Opti-MEM medium (Thermo Fisher Scientific, Waltham, USA) to complete the reaction at a final volume of 300 μl. A second mixture was prepared simultaneously containing 8 μl of lipofectamine LTX and 292 μl of Opti MEM medium. These two reactions were incubated for 5 minutes at room temperature before being mixed and incubated for 30 minutes at room temperature. Meanwhile, HCT 116 cells were washed once with 1X PBS followed by the addition of 2 ml of DMEM culture medium (Thermo Fisher Scientific, Waltham, USA) containing 10% FBS (Wisent, St. John the Baptist, Canada). The selection was started 24 hours later, by adding the antibiotics blasticidine S (5 μg / ml) and hygromycin B (100 μg / ml). The clonal population was maintained over time by the sustained use of these antibiotics, and the induction of constructs was achieved via the addition of 2.5 μg/ml doxycycline (Clontech Laboratories, Mountain View, USA) in the medium at the desired moment.

### EMSA

Oligonucleotides containing the HNF4α consensus DR1 response element (DR1 WT) or a mutated sequence (DR1 MUT) labeled with a biotin molecule at the 5’end (Table 6), were obtained from Integrated DNA Technologies (IDT, San Jose, USA). Hybridized probes were diluted to a concentration of 20 nM for binding reactions. Nuclear extracts were isolated from 293T cell lines transfected with the different pcDNA-DEST47-HNF4α (1-12) plasmids. The cells, cultured at an approximate 90% confluence in 60 mm Petri dishes, were trypsinized and subsequently centrifuged at 1500 × g for 5 minutes. The cell pellet was washed twice with 1 ml of 1X PBS, followed by centrifugation after each wash. The cells were then resuspended in 1 ml of nuclear extraction buffer A (10 mM HEPES pH 7.9, 10 mM KCl, 1.5 mM MgCl 2, 0.34 M sucrose, 10% glycerol, 1 mM DTT, and EDTA-free Protease Inhibitor Cocktail (Roche Applied Science, Penzberg, Germany)). A volume of 10 μl of 10% Triton X-100 solution was added to the cell suspension to obtain a final concentration of 0.1% Triton X-100. The cells were lysed in this buffer for 8 minutes on ice, so as to only rupture the plasma membrane while maintaining the integrity of the nuclear membrane. The cells were centrifuged at 1300 × g for 5 minutes at 4°C. The supernatant, corresponding to the cytoplasmic protein fraction, was collected and stored at −80°C. The nuclei-containing pellets were next resuspended in 150 μl of nuclear extraction buffer B (1% Triton X-100, 150 mM NaCl, 20 mM Tris pH 7.4, 1 mM DTT, and cOmplete EDTA-free Protease Inhibitor Cocktail). The nuclear fraction was incubated with stirring for 30 to 60 minutes at 4°C, before being stored at −80 °C. The nuclear and cytoplasmic fractions were centrifuged at 20,000 × g for 15 minutes to eliminate cell debris. These fractions were assayed using the Pierce BCA Protein Assay kit (Thermo Fisher Scientific, Waltham, USA) as recommended by the manufacturer. Immunoblots of the HNF4α, histone H3 and GAPDH proteins were performed in order to validate the quality of the nuclear extraction.

The binding reactions were prepared in several incubation steps at room temperature, in a final volume of 20 μl. A first incubation of 10 minutes was carried out by mixing 6 μl of nuclear extracts (6 μg), 2 μl of 10X binding buffer (100 mM Tris pH 7.5, 10 mM EDTA, 1 M KCl, 50% glycerol and 1 mM DTT) and 1 μl poly [d (IC)] (1 μg/μl) (cat. # 10108812001, Roche Applied Science, Penzberg, Germany). The biotinylated probe was then added to the mixture at a final concentration of 1 nM and incubated for 25 minutes. Supershift was performed by adding 1.5 μl of purified GFP monoclonal antibody (0.4 μg/μl) (cat. # 11814460001, Roche Applied Science, Penzberg, Germany) for an additional incubation period of 15 minutes. A 2 μl volume of 6X-EMSA loading buffer (6X TBE, 30% glycerol, 0.125% bromophenol blue) was subsequently added to the reaction mixture prior to loading on a non-denaturing polyacrylamide gel.

The non-denaturing polyacrylamide gels were prepared using the SureCast Gel HandCast System (Invitrogen, Carlsbad, USA), in a “mini gel” format (8 × 8 × 0.1cm) at a concentration of 4 % acrylamide: bis, 10% glycerol and TBE 0.5X (45 μM Tris, 45 μM boric acid, 1 mM EDTA pH 8.0). A 60-min pre-migration at 100 V was performed using cold 0.5X TBE buffer as migration buffer. The samples were then loaded onto the gel and migrated at 100 V for 100 minutes. Binding reactions were transferred onto a positively-charged nylon membrane Amersham Hybond-N + (cat #RPN203B, GE Healthcare Life Sciences, Marlborough, USA). The blot was performed in 0.5X TBE buffer at 200 mA for 90 minutes at 4°C. Membrane crosslinking was subsequently achieved using a UV Stratalinker 2400 (Stratagene, San Diego, USA), equipped with UV light bulbs (254 nm, 5 × 15 W). The auto-crosslink function was used, equivalent to an emitted energy of 1200 μJ (x 100) for 45 seconds.

The steps leading to detection of the probe on the positively-charged nylon membrane were performed using the Chemiluminescent Nucleic Acid Detection Module (Thermo Fisher Scientific, Waltham, USA). Detection of biotinylated probes was performed using the ChemiDoc MP imaging system (Bio-Rad, Hercules, USA) and Amersham Hyperfilm ECL autoradiographic films (GE Healthcare Life Sciences, Marlborough, USA).

### Quantitative real-time PCR

The transactivation of different HNF4α target genes by its isoforms was analyzed by real-time quantitative PCR (RT-qPCR), using LightCycler 96 (Roche Applied Science, Penzberg, Germany) and cDNA from 48hr-induced HCT 116 HNF4α (1-12) -GFP lines. The oligonucleotides used for the amplification of the tested genes were obtained from Integrated DNA Technologies (IDT, San Jose, USA) (Supplementary Table 7). The reactions were performed with the SYBR Green Master Fast Start reaction mixture (Roche Applied Science, Penzberg, Germany) as recommended by the manufacturer. Analysis of the amplification and melting curves was performed using LightCycler 96 software version 1.1.0.1320 (Roche Applied Science, Penzberg, Germany). The relative expression of the genes was calculated by comparison with the TBP reference gene with the formula E_target_ (Cp_Control_-Cp_Sample_) X E_reference_ (Cp_Sample_-Cp_Control_).

### Transcriptomics

#### Preparation of total RNA

The HCT 116 HNF4α (1-12) -GFP and HCT 116 Flp-In T-Rex lines were inoculated in 60 mm Petri dishes and incubated for 48 hours in the presence of 2.5 μg/ml doxycycline. Total RNA of these 13 lines was extracted into triplicates using RNeasy RNA isolation kit (QIAGEN). The concentration as well as the quality of the RNAs was evaluated first by NanoDrop (Thermo Fisher Scientific, Waltham, United States) via the RNomics platform of the Université de Sherbrooke and by a 2100 Bioanalyzer (Agilent, Santa Clara, USA). The samples were then sent to McGill University’s Innovation Center and Génome Québec.

#### Preparation of libraries and sequencing

Libraries were generated from 250 ng of total RNA. Enrichment of the mRNA was performed using the NEBNext Poly (A) Magnetic Insulation Module (New England Biolabs, Ipswich, USA). The cDNA synthesis was performed via the use of NEBNext RNA First Strand Synthesis and NEBNext Ultra Directional RNA Second Strand Synthesis modules (New England Biolabs, Ipswich, USA). The final steps for the preparation of the libraries were carried out using the NEBNext Ultra II DNA Library Prep Kit for Illumina (New England Biolabs, Ipswich, USA). PCR adapters and primers were obtained from New England Biolabs. The libraries were quantified using Quant-iTT PicoGreen® dsDNA Assay Kit (Thermo Fisher Scientific, Waltham, USA) and Kapa Illumina GA with Revised Primers-SYBR Universal Fast Kit (Kapa Biosystems, Wilmington, USA). The average size of the RNA fragments was determined using the LabChip GX instrument (PerkinElmer, Waltham, USA). Sequencing of the libraries was performed using NovaSeq 6000 (Illumina, San Diego, USA) using a S2 PE100 protocol.

### Sequence alignment

The quality of the sequencing results was visualized using the FastQC 0.11.5 tool (Wingett and Andrews, 2018). The sorting of the data according to their quality score was performed using the Trimmomatic 0.36 software (Bolger et al., 2014). The transcriptome used for the alignment of the reads was constructed from the human genome GRCh38.p12 and annotations from RefSeq (NCBI Homo sapiens Annotation Release 109) (O’Leary et al., 2016). Transcriptome alignment and quantification were performed using the Kallisto 0.44.0 tool (Bray et al., 2016). Results from Kallisto counts were used to quantify transcripts, as well as genes by addition of transcripts. The DESeq2 1.14.1 software was used to calculate the differential expression of transcripts and genes for each isoform in comparison with the control sample (Love et al., 2014).

### Mass spectrometry

#### Cell culture and induction of different constructions

The stable HCT 116 HNF4α (1-12) -BioID2-3Xmyc and HCT 116 BioID2-3Xmyc-control cell lines were cultured in three different SILAC media designated as light (R0K0), medium (R6K4) and heavy (R10K8). SILAC media contained DMEM + 4.5 g/L glucose, L-glutamine, sodium pyruvate (Thermo Fisher Scientific, Waltham, USA) supplemented with 10% of triple-dialyzed FBS (Thermo Fisher Scientific, Waltham, USA), 10 mM HEPES, 2 mM GlutaMAX, 100 U/ml penicillin and 100 μg/ml streptomycin. To these media were added different L-arginine and L-lysine isotopes at final concentrations of 42 μg/ml and 63.5 μg/ml respectively to obtain light, medium and heavy media. The light medium contained L-arginine R0 (Sigma-Aldrich A6969, St. Louis, USA) and L-lysine K0 (Sigma Aldrich A8662, St. Louis, USA). The medium medium contained L-arginine R6 (Cambridge Isotope Laboratories, Inc. CLM-2265, Tewksbury, USA) and L-lysine K4 (Cambridge Isotope Laboratories, Inc. DLM-2640, Tewksbury, USA). The heavy medium contained L-arginine R10 (Cambridge Isotope Laboratories, Inc. CNLM-539, Tewksbury, USA) and L-lysine K8 (Cambridge Isotope Laboratories, Inc. CNLM-291, Tewksbury, USA). The different SILAC culture media were then filtered on a Stericup filtration unit (EMD Millipore, Burlington, USA) prior to use. The HCT 116 BioID2-3Xmyc cells, serving as a control condition for the mass spectrometry experiments, were cultured in the light medium. The HCT 116 HNF4α (1-6) -BioID2-3Xmyc lines were cultured in the medium medium while the HCT 116 HNF4α (7-12) -BioID2-3Xmyc lines were cultured in the heavy medium. Culture in SILAC medium involved a minimum of 5 passages at a ratio of 1:4 followed by incubation for 2-3 days in polystyrene Petri dishes 100 or 150 mm, at 37°C in a controlled atmosphere at 5% CO2.

A 150 mm Petri dish was used for each condition. During the last passage in SILAC medium, doxycycline (Clontech Laboratories, Mountain View, USA) was added to the cells at a concentration of 2.5 μg/ml 48 hours before the streptavidin pulldown experiment, when the cells were at a confluence of 40 to 50%. The next day, 24 hours prior to the pulldown, biotin (Sigma-Aldrich, St. Louis, USA) was added to the cells at a final concentration of 50 μM. The cells were then washed three times with 1X PBS, trypsinized and centrifuged at 1500 × g for 5 minutes at 4°C. The cell pellets were washed again twice with 1 × PBS.

#### Immunoprecipitation of biotinylated proteins

The cell pellets were lysed in 1 ml of RIPA buffer (50 mM Tris-HCl, pH 7.5, 150 mM NaCl, 1.5 mM MgCl 2, 0.1% SDS, 1% IGEPAL CA-630 (Sigma-Aldrich, St. Louis, USA)), 1 mM PMSF, 0.4% sodium deoxycholate, 1 mM DTT, and the EDTA-free Protease Inhibitor Cocktail inhibitor per 150 mm. The cell lysates were rotated for 20 to 30 minutes at 4°C and sonicated on ice with a Sonic Dismembrator Model 120 (Thermo Fisher Scientific, Waltham, USA) at 30% amplitude three times for 10 to 15 seconds interspaced by 5-sec pauses. A 40 μl aliquot of each sample was taken for assay using the Pierce BCA Protein Assay kit (Thermo Fisher Scientific, Waltham, USA). The samples were then rotated for an additional hour at 4°C, after the addition of 10 μl of 0.1 M EGTA (1 mM final). A combination of the three SILAC conditions (light, medium and heavy) was performed prior to the pulldown by mixing 3.5 mg of total extract of each condition and supplementing to a volume of 3 ml with RIPA buffer. A 45 μl aliquot of 20% SDS was added to each sample (0.4% final). The samples were rotated for 15 minutes at 4°C and subsequently centrifuged at 20,000 × g for 20 minutes at 4°C.

High performance streptavidin Sepharose beads (GE Healthcare Life Sciences, Marlborough, USA) were used for the pulldown at 50 μl per combined sample. The beads were washed three times with 1 ml of RIPA buffer containing 0.4% SDS. Washings were performed by rotating the beads for 5 minutes at 4°C and subsequently centrifuging at 6000 × g for 3 minutes at 4°C before removing the supernatant. The washed beads were resuspended at 50% concentration in the same buffer and added to the samples in 5 ml tubes. The samples were rotated for 3 hours at 4°C. The beads were centrifuged at 6000 × g for 3 minutes at 4°C, and a 100 μl aliquot of the supernatant was removed. The beads were transferred to Low Binding microtubes (Sarstedt, Nümbrecht, Germany) and washed according to the parameters described above. A first wash was performed with 1.5 ml of BioID wash buffer (2% SDS, 50 mM Tris-HCl, pH 7.5), followed by three washes with 1 ml of RIPA buffer containing 0.4% SDS and five washes with 1 ml of 20 mM ammonium bicarbonate buffer. An aliquot corresponding to 5% of the total amount of beads was collected. The final washed beads were stored at −80°C, as were all the aliquots collected during the experiment.

#### Reduction, alkylation and digestion of proteins

All buffers used in this stage were prepared with MS-grade water. The protein reduction step was carried out by incubating the beads in 100 μl of 20 mM ammonium bicarbonate buffer containing 10 mM DTT (Thermo Fisher Scientific, Waltham, USA) with stirring (1250 rpm) for 30 minutes at 60°C. The alkylation of the proteins was carried out by adding another 100 μl of 20 mM ammonium bicarbonate buffer, and then supplementing with 15 mM iodoacetamide (Sigma-Aldrich, Saint-Louis, United States) final before stirring for 1 hour at room temperature away from light. The IAA was then neutralized by completing with 15 mM DTT and stirring for 10 minutes at 37°C. The proteins were digested by adding 1 μg Pierce MS-grade trypsin (Thermo Fisher Scientific, Waltham, USA) and incubated overnight at 37°C with shaking.

#### Purification and desalting of the peptides on C18 columns

Digestion was stopped by adding 1% formic acid (FA) (Thermo Fisher Scientific, Waltham, USA) to a total volume of 200 μl, followed by stirring for 5 minutes at room temperature. The beads were spun at 6000 × g for 3 minutes before harvesting the supernatant and transferring to a new low binding microtube. The beads were resuspended in 200 µl of buffer containing 60% acetonitrile (ACN) (Sigma-Aldrich, St. Louis, USA) and 0.1% FA, and subsequently stirred for 5 minutes at room temperature. The supernatant was harvested and combined with that obtained previously. These samples were thereafter concentrated by a centrifugal evaporator at 65°C until complete drying (approximately 2 hours), and resuspended in 30 μl of 0.1% trifluoroacetic acid (TFA) buffer (Sigma-Aldrich, St. Louis, USA). The peptides were purified with ZipTip 10-μl micropipette tips containing a C18 column (EMD Millipore, Burlington, USA). The ZipTip was first moistened by suctioning 10 μl of 100% ACN solution three times, then equilibrated by suctioning 10 μl of 0.1% TFA buffer three times. Each peptide sample was passed on the balanced ZipTip by 10 succeeding up-and-downs of 10 μl of the sample. This step was performed three times in order to pass the entire sample on the column. The ZipTip was then washed with 10 μl of 0.1% TFA buffer three times. The elution of the peptides was performed in a new low-binding microtube, 10 times with a volume of 10 μl of 50% ACN and 0.1% FA buffer. This step was carried out three times to obtain a final volume of 30 μl. The peptides were then concentrated by centrifugal evaporator at 65°C until complete drying (approximately 30 minutes) and then resuspended in 25 μl of 1% FA buffer. Peptides were assayed using a NanoDrop spectrophotometer (Thermo Fisher Scientific, Waltham, USA) and read at an absorbance of 205 nm. The peptides were then transferred to a glass vial (Thermo Fisher Scientific, Waltham, USA) and stored at −20°C until analysis by mass spectrometry.

#### LC-MS/MS analysis

Trypsin-digested peptides were separated using a Dionex Ultimate 3000 nanoHPLC system. Ten μl of sample (a total of 2 μg) in 1% (vol/vol) formic acid were loaded with a constant flow of 4 μl/min onto an Acclaim PepMap100 C18 column (0.3 mm id × 5 mm, Dionex Corporation). After trap enrichment, peptides were eluted onto an EasySpray PepMap C18 nano column (75 μm × 50 cm, Dionex Corporation) with a linear gradient of 5-35% solvent B (90% acetonitrile with 0.1% formic acid) over 240 minutes with a constant flow of 200 nl/min. The HPLC system was coupled to an OrbiTrap QExactive mass spectrometer (Thermo Fisher Scientific Inc) via an EasySpray source. The spray voltage was set to 2.0 kV and the temperature of the column set to 40°C. Full scan MS survey spectra (m/z 350-1600) in profile mode were acquired in the Orbitrap with a resolution of 70,000 after accumulation of 1,000,000 ions. The ten most intense peptide ions from the preview scan in the Orbitrap were fragmented by collision-induced dissociation (normalized collision energy 35% and resolution of 17,500) after the accumulation of 50,000 ions. Maximal filling times were 250 ms for the full scans and 60 ms for the MS/MS scans. Precursor ion charge state screening was enabled and all unassigned charge states as well as singly, 7 and 8 charged species were rejected. The dynamic exclusion list was restricted to a maximum of 500 entries with a maximum retention period of 40 seconds and a relative mass window of 10 ppm. The lock mass option was enabled for survey scans to improve mass accuracy. Data were acquired using the Xcalibur software.

#### Protein identification by MaxQuant analysis

The raw files were analyzed using the MaxQuant version 1.6.2.2 software (Cox and Mann, 2008) and the Uniprot human database (16/07/2013). Isoform analyses were initially performed separately in order to obtain enrichment ratios for the complete sequence of each isoform. A common analysis was then performed to integrate all raw MS/MS analysis files. The MaxQuant software default settings were used, except for the following parameters: multiplicity of 3 SILAC media (R0K0, R6K4 and R10K8), identification values “PSM FDR”, “Protein FDR” and “Site decoy fraction” 0.05, minimum ratio count of 1 and selection of the “Re-quantify” option. Following the analysis, the results were sorted according to several parameters. Proteins positive for at least one of the “Reverse”, “Only.identified.by.site” and “Potential.contaminant” categories were eliminated, as well as proteins identified from a single peptide. The ratios identified in only one of the three replicas for each experiment were eliminated. The ratios identified in two of the three replicas were eliminated when they were considered to be too divergent, i.e. when the standard deviation was greater than the average of the two enrichment ratios. Outliers for the ratios measured in the three replicas were detected using the Grubbs test at a value of α = 0.05 and then eliminated. Following this sorting, proteins for which no ratio was calculated in at least one experiment were removed. Gene ontology enrichment analyses were performed using the Panther 13.1 tool (Mi et al., 2010). Heatmap visualizations were created using the Morpheus software (https://software.broadinstitute.org/morpheus/).

## DATA DEPOSITION

The HNF4α RNAseq dataset was deposited to the NCBI Gene Expression Omnibus (GEO; https://www.ncbi.nlm.nih.gov/geo/) under the accession number GSE125852.

The mass spectrometry proteomics data was deposited to the ProteomeXchange Consortium via the PRIDE (Vizcaino et al., 2016) partner repository with the dataset identifier PXD012146.

## SUPPLEMENTARY INFORMATION

Supplementary Information includes eight figures and seven tables.

## ACKNOWLEDGEMENTS

This work was supported by the Canadian Institutes of Health Research (CIHR) (grant number MOP-123469 to F.M.B and grant number PJT-156180 to F.B.), the Natural Sciences and Engineering Research Council of Canada (NSERC) (grant number RGPIN-2017-06096 to F.B.) and by the Cancer Research Society (to F.M.B.). F.M.B. and M.S.S. are FRQS Junior II scholars (award number 32956 to F.M.B and 34877 to M.S.). E.L. is recipient of NSERC and FRQS scholarships and J.S. is a recipient of FRQNT and NSERC scholarships. M.S.S., F.B. and F.M.B. are members of the FRQS-funded " Centre de Recherche du CHUS».

## AUTHOR CONTRIBUTIONS

E.L. and J.-P.B. performed the majority of the experiments and analyzed the data. J.S. and M.S. performed the analysis of the RNAseq which includes Figure 5 and Suppl. Figure 6. D.L. provided help with the cloning, the generation of the plasmids and the mass spectrometry analysis. E.J. contributed to the EMSA, as well as the validations by co-immunoprecipitation assays. F.B. and F-M.B. contributed to the design of the experiments, supervised the project and prepared the manuscript.

## Supplementary Figure Legends

**Supplementary Figure 1:**
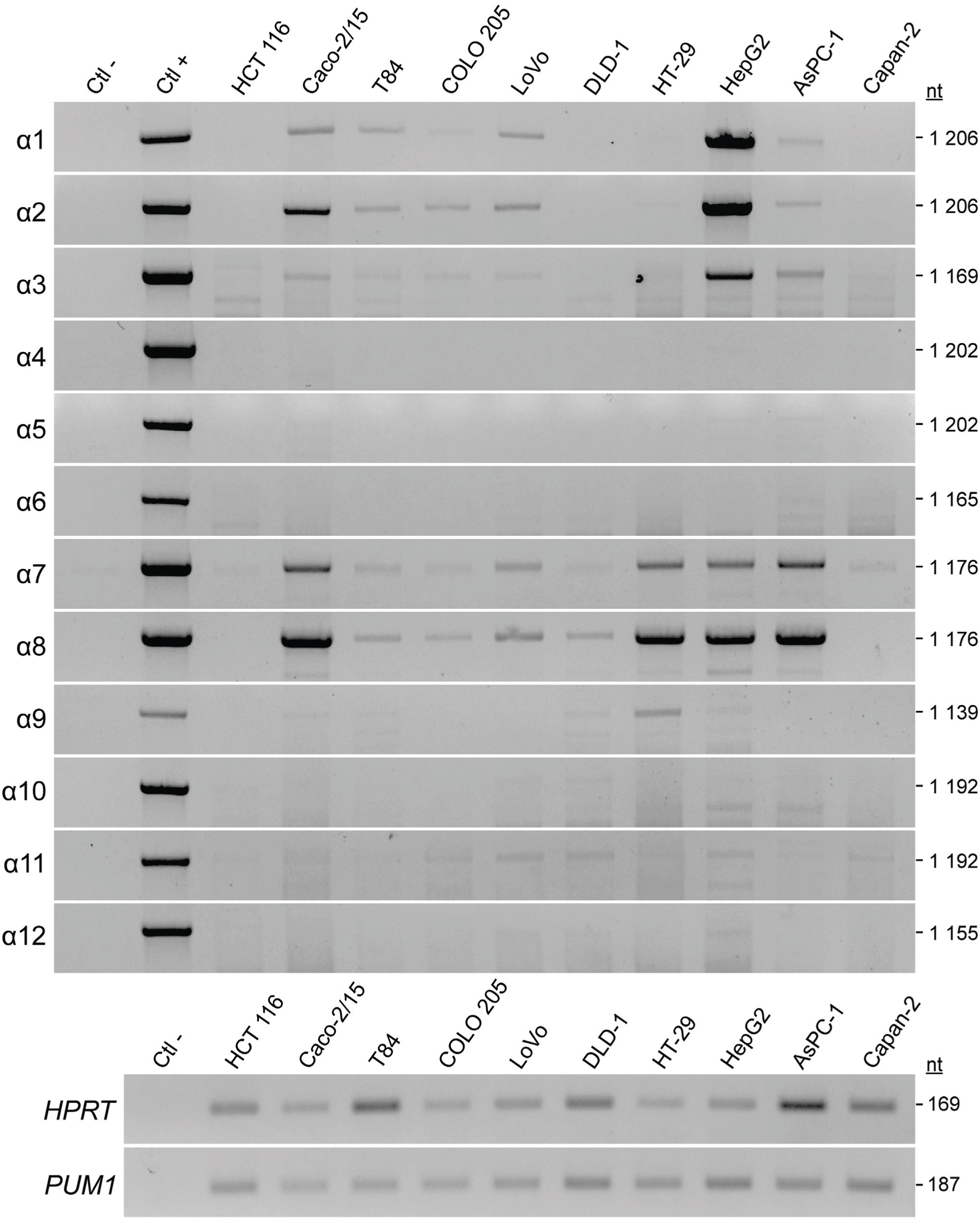
The isoforms of HNF4α isoforms are expressed at varying levels in different human cancer cell lines. The expression of the isoforms was evaluated by semi-quantitative RT-PCR in the indicated cell lines. HPRT and PUM1 genes were used as reference genes.

**Supplementary Figure 2:**
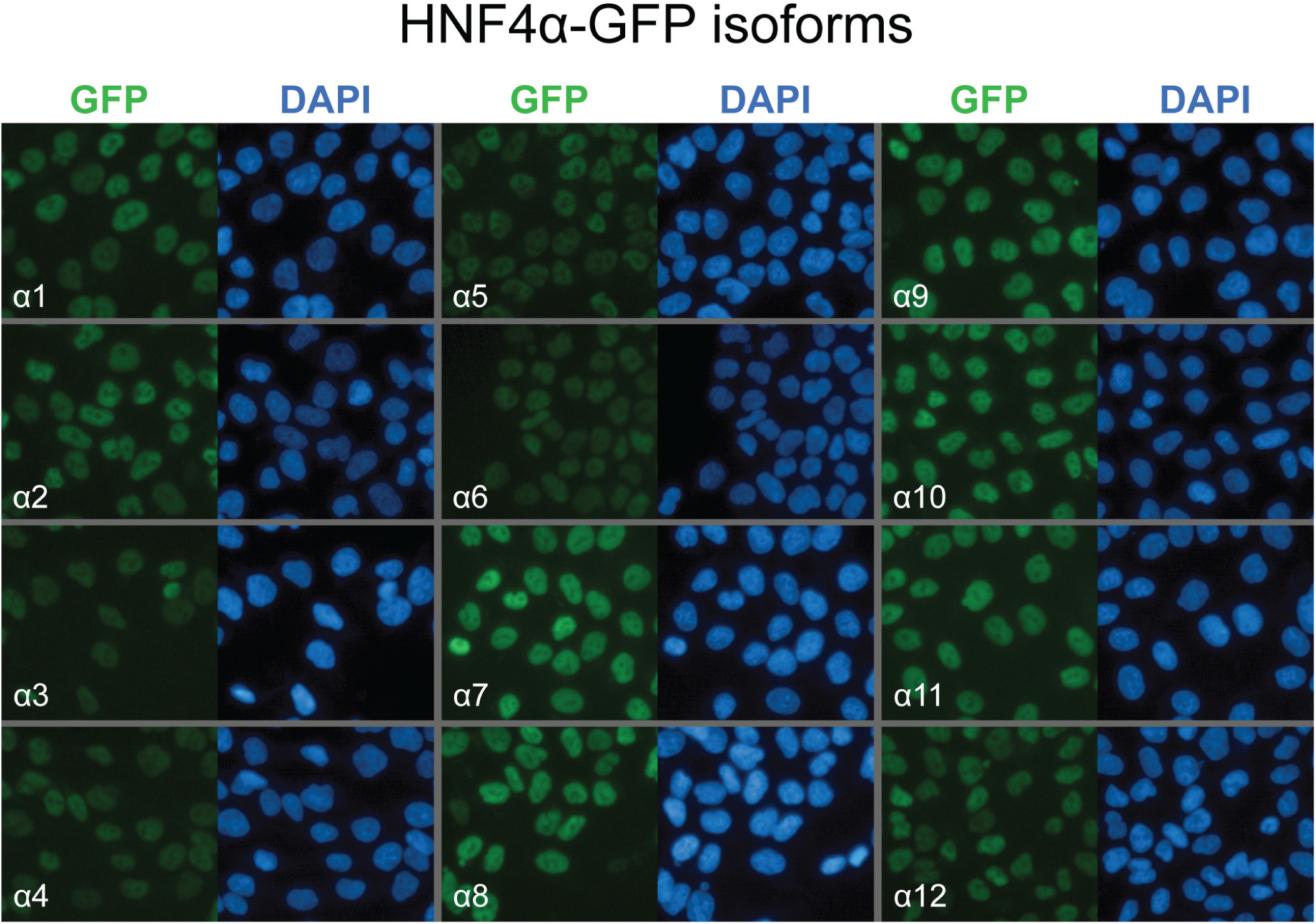
GFP-tagged HNF4α localization by immunofluorescence microscopy. Cells expressing the different GFP-tagged HNF4α isoforms were fixed, labeled with a GFP antibody (green) and the nuclei stained with DAPI (blue).

**Supplementary Figure 3:**
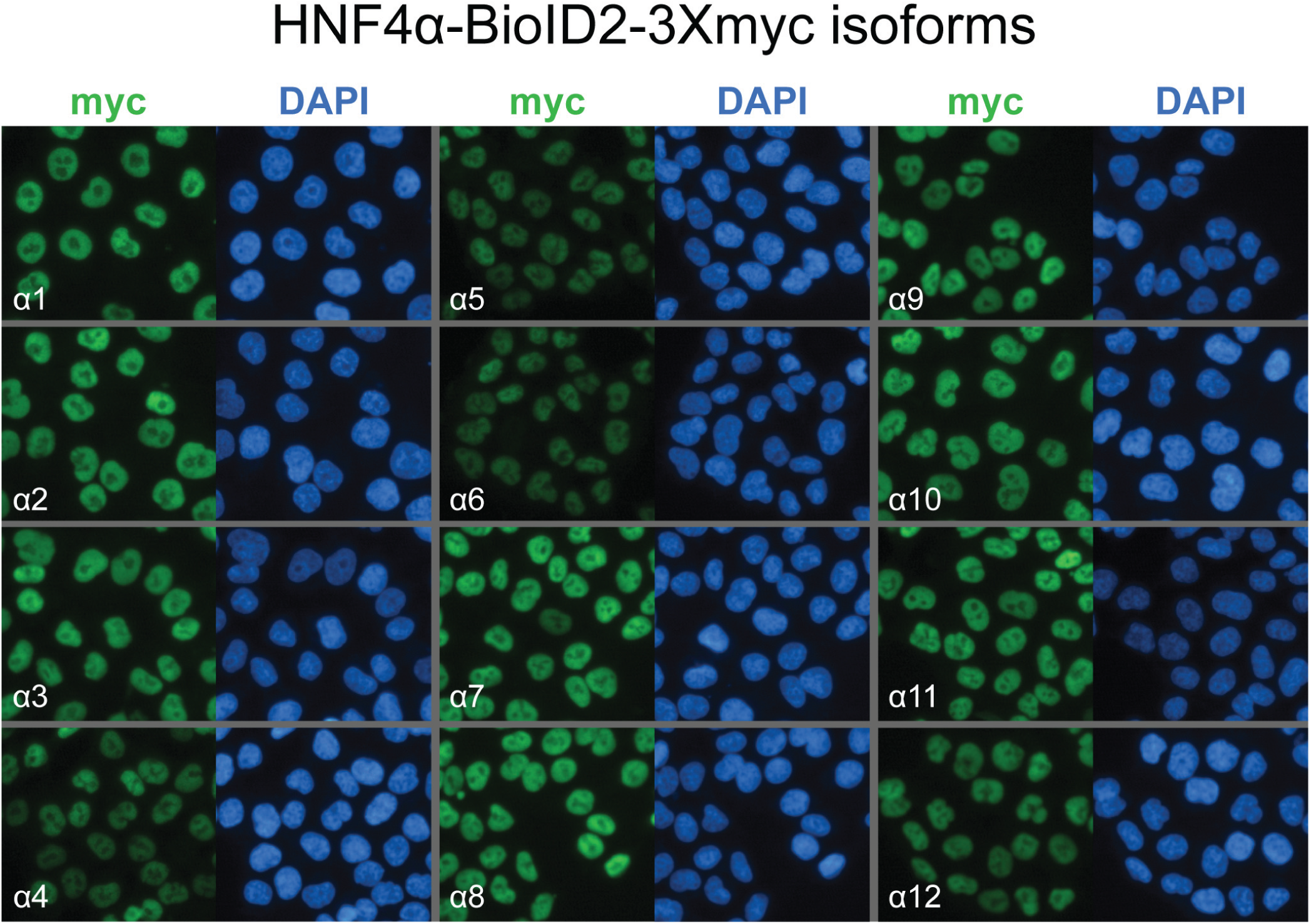
BioID2-3myc-tagged HNF4α localization by immunofluorescence microscopy. Cells expressing the different BioID2-3myc-tagged HNF4α isoforms were fixed, labeled with a myc antibody (green) and the nuclei stained with DAPI (blue).

**Supplementary Figure 4:**
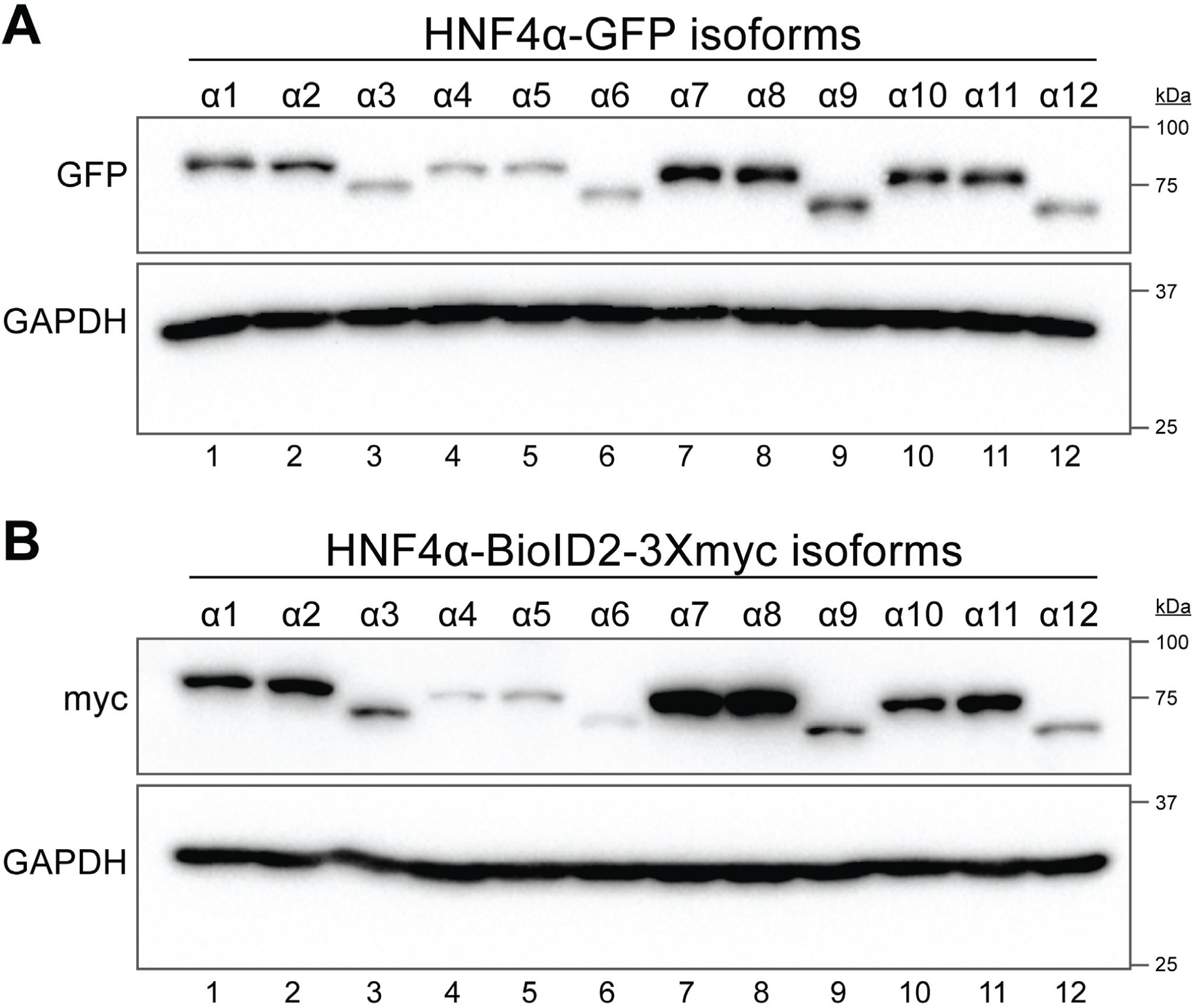
Protein expression analysis of all 12 stable cell lines (GFP and BioID2) by immunoblotting. Protein expression from whole cell lysates of cell lines expressing each of the HNF4α isoforms following induction with doxycycline was assessed by immunoblotting with a GFP antibody (A) and a myc antibody (B).

**Supplementary Figure 5:**
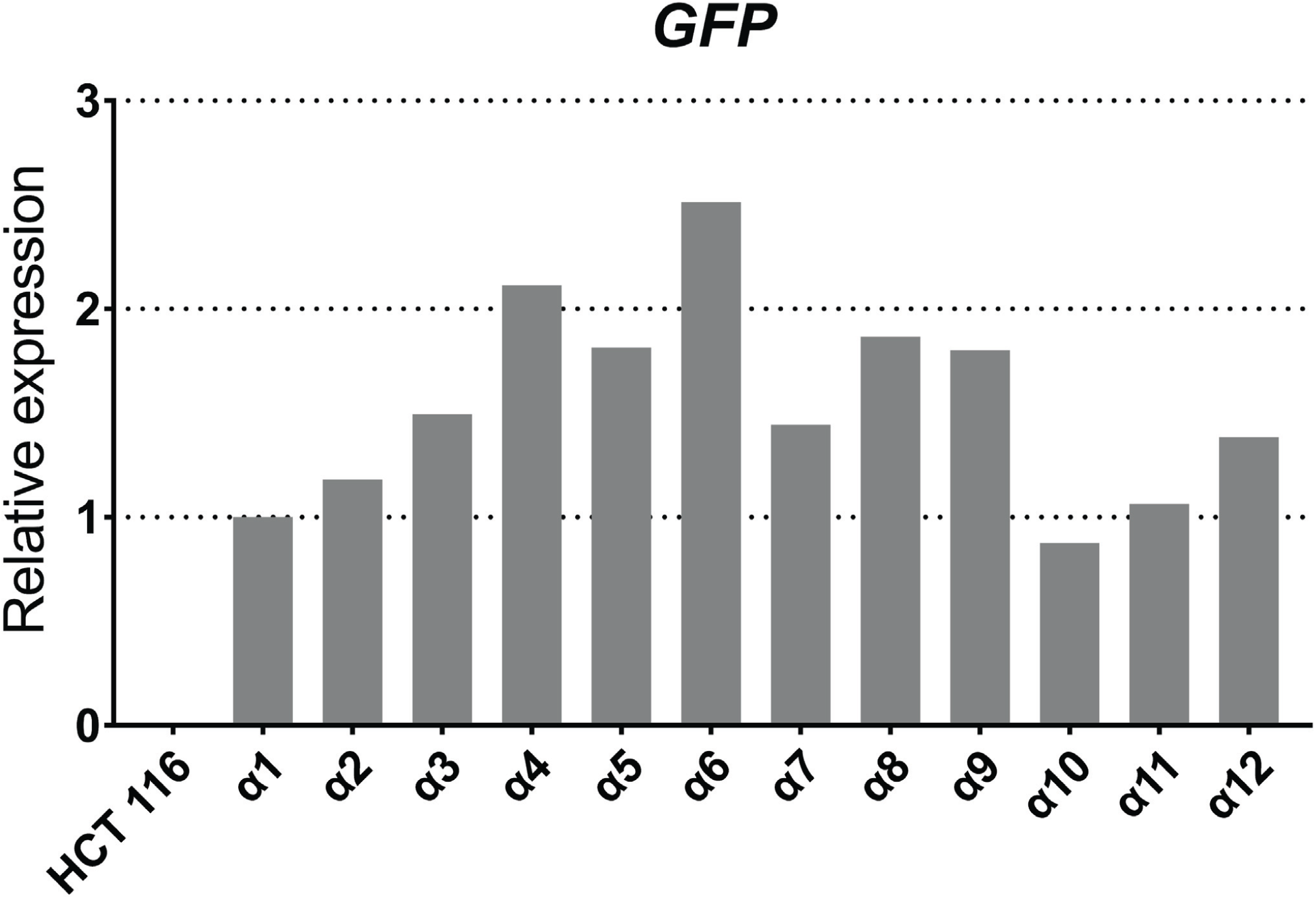
Relative mRNA expression of HNF4α isoforms in the stable cell lines. Analysis of the transcript levels in each of the cell lines expressing the GFP-tagged HNF4α isoforms by qPCR using oligonucleotides recognizing GFP.

**Supplementary Figure 6:**
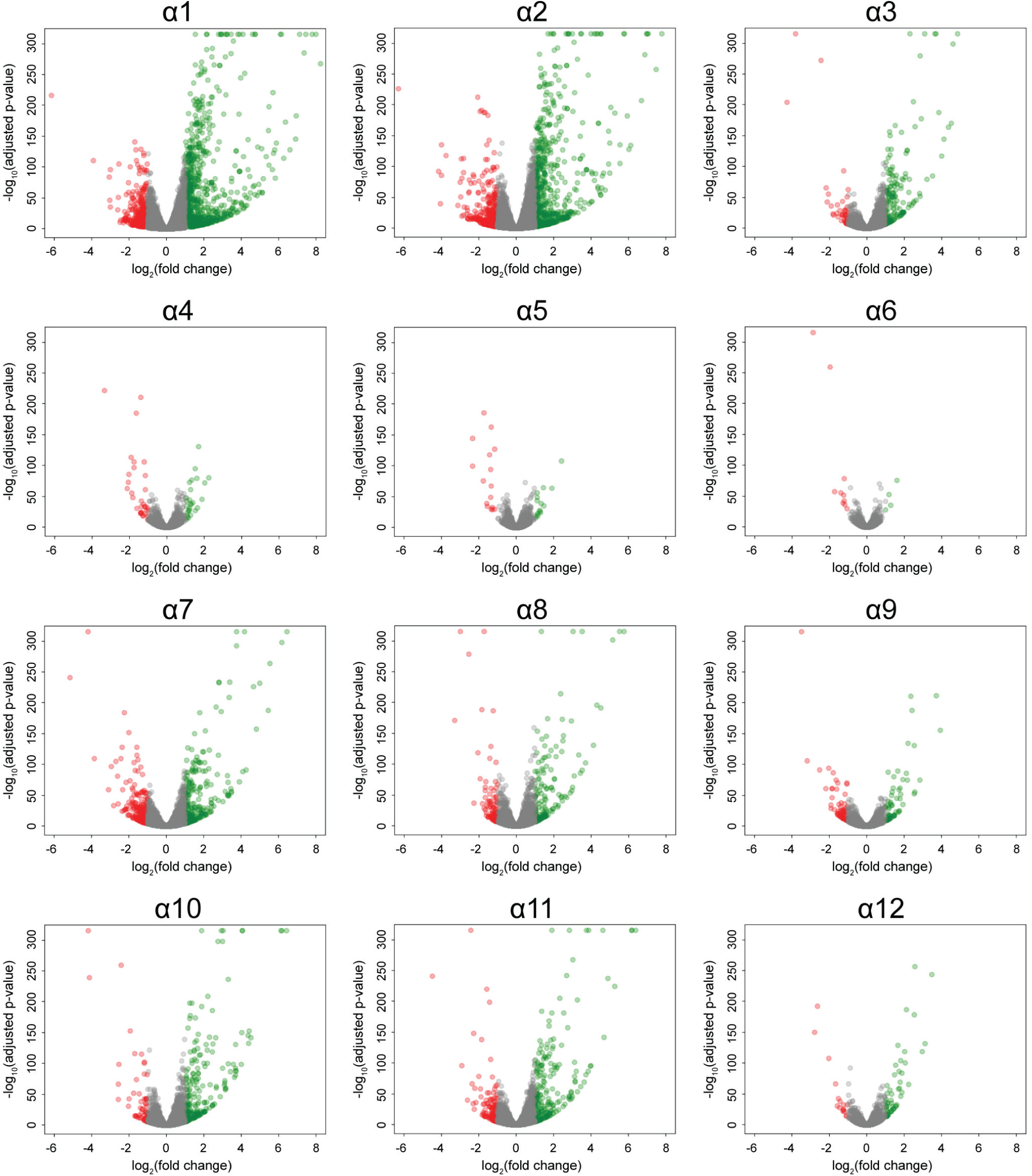
Volcano plots of statistical significance against fold-change comparing each HNF4α isoforms with the control. The base 10 logarithm of the adjusted p-value is presented as a function of the logarithm in base 2 of the modulation level (FoldChange). A minimum absolute modulation threshold of 2 combined with an adjusted p-value threshold ≤ 0.001 was used. Genes above this threshold are modulated upwards (in green) or downward (in red).

**Supplementary Figure 7:**
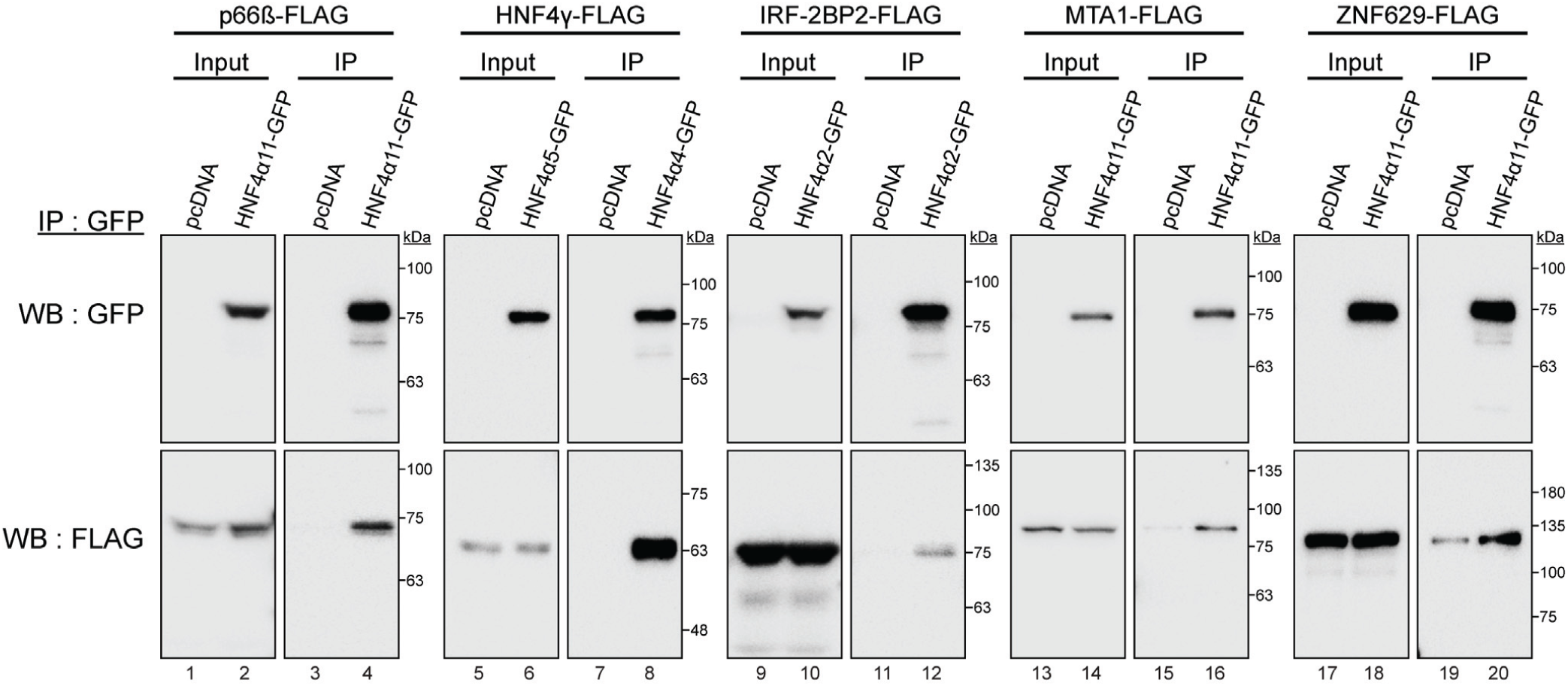
Validation of the interactions between the HNF4α isoforms and five proteins identified by the BioID approach. 293T cells were co-transfected for 48 hours with the indicated GFP-tagged HNF4α isoform and potential FLAG-tagged interaction partners (GATAD2B, HNF4γ, IRF-2BP2, MTA1 or ZNF629). Protein complexes were immunoprecipitated using a GFP antibody from total cell extracts. The co-immunoprecipitations were revealed with GFP and FLAG antibodies, making it possible to validate the interaction between HNF4α and the five proteins indicated above (n = 2).

**Supplementary figure 8:**
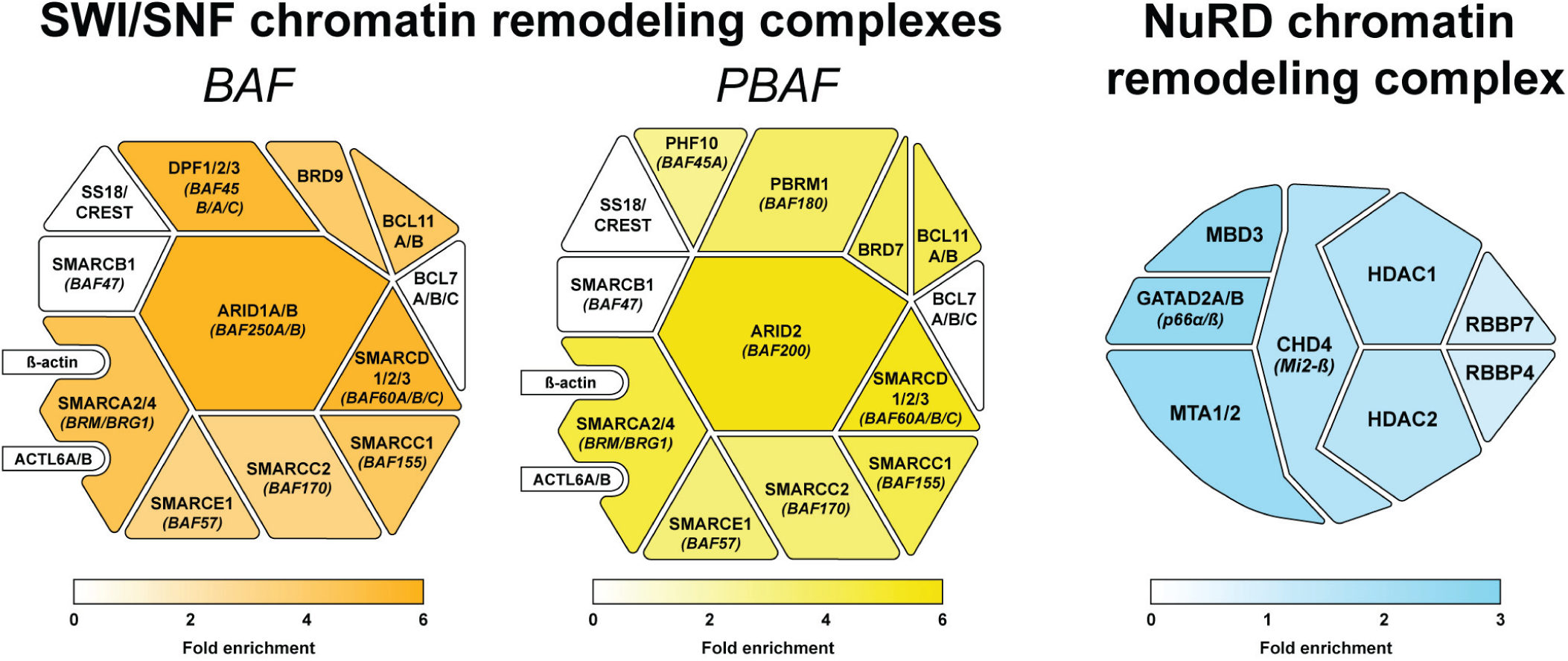
Chromatin remodeling complex identification through BioID-MS. The three main complexes with the associated subunits are displayed with the average standardized SILAC enrichment scores for BAF / PBAF and NuRD complexes.

**Supplementary Table 1:**
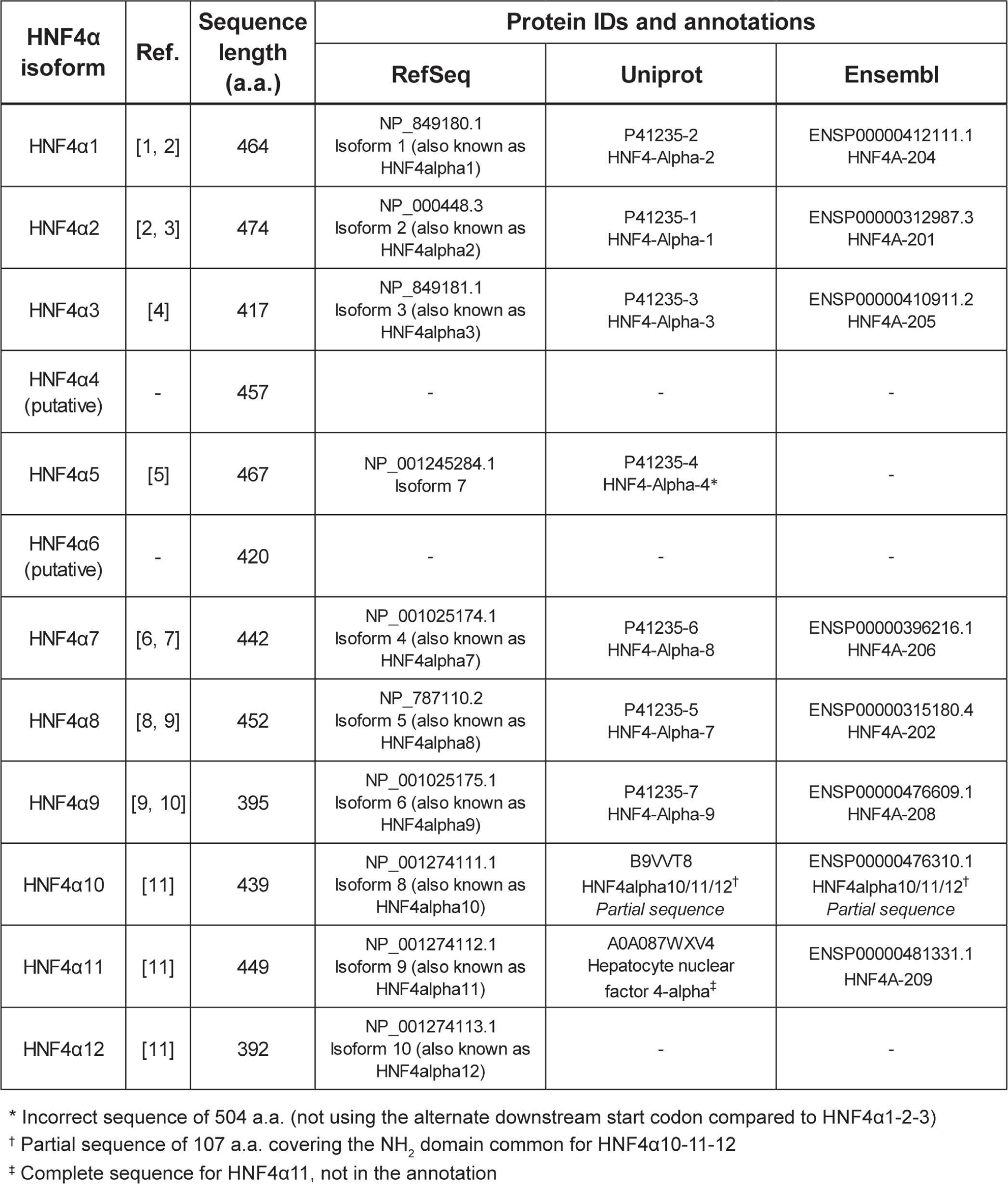
Identification and annotations of HNF4α isoforms in RefSeq, Uniprot and Ensembl.

**Supplementary Table 2:** RNAseq results (gene name, intensities, fold change, p-values).

**Supplementary Table 3:** RNAseq quality control statistics.

**Supplementary Table 4:** Listing of the different SILAC label combinations.

| #<br>sample | Light (R0K0) | Medium (R6K4) | Heavy (R10K8) |
| --- | --- | --- | --- |
| | HCT 116<br>BioID2 | HCT 116<br>P1-HNF4α-<br>BioID2-3Xmyc | HCT 116<br>P2-HNF4α-<br>BioID2-3Xmyc |
| 1 | Negative control | HNF4α1 | HNF4α7 |
| 2 | Negative control | HNF4α2 | HNF4α8 |
| 3 | Negative control | HNF4α3 | HNF4α9 |
| 4 | Negative control | HNF4α5 | HNF4α10 |
| 5 | Negative control | HNF4α4 | HNF4α11 |
| 6 | Negative control | HNF4α6 | HNF4α12 |

**Supplementary Table 5:** Mass spectrometry data for each of the HNF4α isoforms.

**Supplementary Table 6:**
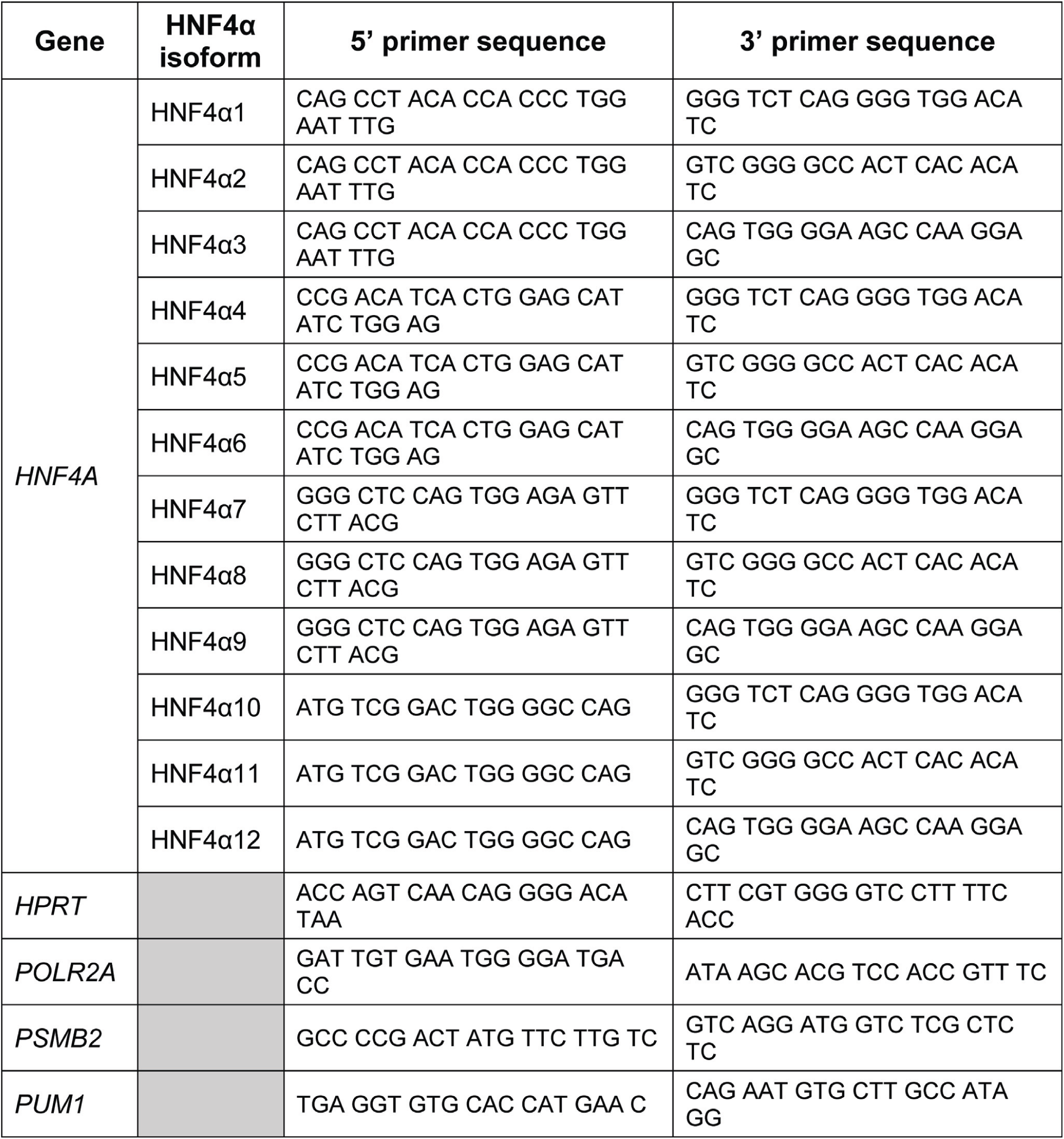
Semi-quantitative RT-PCR primers.

**Supplementary Table 7:**
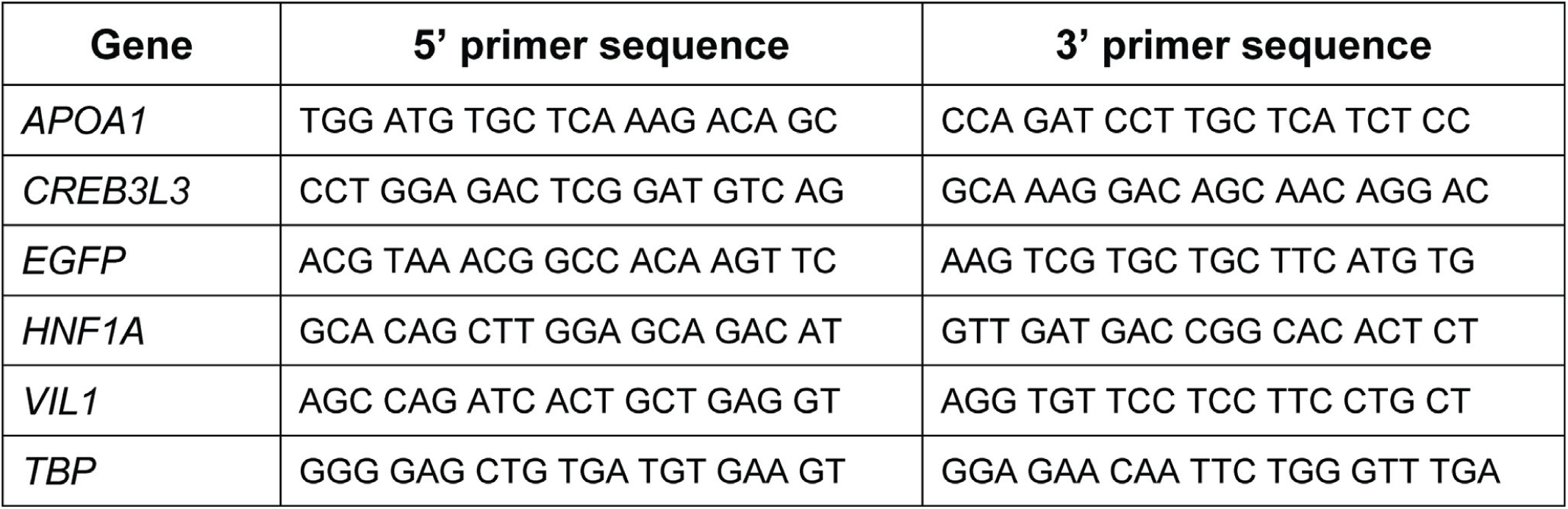
Real-time quantitative RT-PCR primers.

